# Early-Life Wildfire Smoke Exposure Is Associated with Long-Term Systemic Immune Remodeling and Epigenetic Reprogramming

**DOI:** 10.64898/2026.08.27.742220

**Authors:** Cora E. Layman, David W. Morrow, Taylor J. Caron, Kandace Wheeler, Brett A. Davis, Paige Bergstrom, Katinka Vigh-Conrad, Tanner J. Anderson, G.W. McElfresh, Kirstin N. Sterner, Baptiste Sadoughi, Noah Snyder-Mackler, Scott G. Hansen, Benjamin N. Bimber, Christina Lancioni, Lucia Carbone, Mariam Okhovat

## Abstract

Wildfire smoke is an escalating global public health threat exposing millions of people, including children, to hazardous air pollution each year. Although wildfire smoke toxicants have been linked to a range of adverse health outcomes, including immune dysregulation, the long-term consequences of real-world pediatric wildfire smoke exposure on health and development remain largely unknown. To investigate the persistent effects of early-life exposure on immune health, here we leveraged a cohort of rhesus macaques that experienced nine consecutive days of hazardous wildfire smoke exposure in infancy during the 2020 Oregon Labor Day wildfires. By integrating *ex vivo* immune stimulations, multiplex cytokine profiling, single-cell transcriptomics, and genome-wide DNA methylation profiling, we identified persistent immunological consequences across molecular and functional levels. We found that a single severe postnatal exposure, in the first three months of life, was associated with persistent change in the innate immune response, including reduced pro-inflammatory cytokine response to a bacterial endotoxin, with subtle but consistent transcriptional changes in myeloid cells, particularly among males. Wildfire smoke exposure was also associated with changes in proportion of B and T/NK cells, and within the T/NK cell compartment, exposed animals exhibited an expansion of cytotoxic cells. Consistent with this, CD8^+^ T cells displayed extensive transcriptional remodeling and shifted toward more differentiated effector states, with the greatest differentiation observed in animals exposed at the youngest ages. Genome-wide DNA methylation profiling identified smoke-associated methylation changes consistent with acceleration of epigenetic aging, as well as persistent epigenetic alterations impacting genes involved in oxidative stress responses, innate immunity, T cell differentiation, and hematopoiesis. These findings demonstrate that a single severe wildfire smoke exposure during a critical developmental window is associated with extensive immune and epigenetic remodeling that persist years after exposure, providing new insight into the long-term biological consequences of early-life wildfire smoke exposure.

## INTRODUCTION

Wildfire smoke exposure is a major and worsening environmental health concern, exposing millions of people worldwide to hazardous air pollution each year. Because wildfire smoke can travel hundreds to thousands of miles, it can expose entire communities far beyond the immediate fire perimeter. Unlike chronic urban air pollution, wildfire smoke exposure is typically episodic, characterized by discrete periods of acute exposure that can range from hours to several weeks, often with little advance warning, making them difficult to avoid. In the United States alone, an estimated 212 million people lived in counties affected by wildfire smoke in 2011^1^. Over the following decade, the number of days of heavy smoke increased in communities representing 87.3% of the whole US population^2^, and the number of people experiencing at least one day of hazardous-level wildfire smoke annually rose 27-fold, reaching nearly 25 million people in 2020^2^. Children represent one of the largest and most vulnerable populations exposed to wildfire smoke. A 2017 study estimated that approximately 7.4 million children in the United States were exposed annually^3^. Unfortunately, this estimate is likely substantially higher today and is expected to continue rising as populations grow and climate change increases the frequency, duration, and intensity of wildfires^4^. This worsening public health burden underscores the urgent need to understand the biological consequences of wildfire smoke exposure, particularly during early-life, when exposure to environmental toxicants can have lasting effects on development and health^5,6^.

Wildfire smoke is a complex mixture of combustion-derived compounds, including carbon monoxide, nitrogen oxides, volatile organic compounds (VOCs), polycyclic aromatic hydrocarbons (PAHs), heavy metals, and particulate matter. Although many constituents of wildfire smoke contribute to its toxicity^7^, fine particulate matter (PM_2.5_; particles ≤2.5 μm in diameter) is particularly well-characterized and biologically relevant. Due to its small size, PM_2.5_ can readily penetrate deep into the lungs, cross the alveolar barrier, and enter the bloodstream, where it can reach various tissues and cause cellular injury, oxidative stress, inflammation, mitochondrial dysfunction, epigenetic alterations and DNA damage^8^. Accordingly, ambient PM_2.5_, which is often used as a primary metric for quantifying ambient air pollution, has been linked to a broad range of adverse health issues, including death, respiratory and cardiovascular problems, adverse fertility and pregnancy outcomes, impaired neurodevelopment, and immune dysfunction^9,10^.

The immune system is increasingly recognized as a major biological system affected by smoke and PM_2.5_ exposure. These effects are of particular concern during early-life because children receive a greater dose of inhaled pollutants per unit of body weight, and their respiratory and immune systems are still undergoing major development^11^. Consistent with this heightened environmental vulnerability, some studies report associations between pediatric exposure to urban air pollution and tobacco smoke and persistent alteration in immune function and increase susceptibility and severity of immune-related diseases later in life^12–14^. However, many studies have focused on adult populations and on the acute health effects occurring during or immediately following smoke exposure^15^. Thus, we have a limited understanding of the long-term immune impacts of wildfire smoke exposure during early-life, and the biological mechanisms underlying long-term immune dysregulation. Addressing these questions in humans is challenging due to logistical and ethical constraints, particularly considering the potentially long latency between childhood exposure and the emergence of health effects. While *in vitro* systems and rodent models can provide important mechanistic insights, non-human primates models such as the rhesus macaque, more closely recapitulate human physiology, genetics, and development making them better suited to investigating the long-term immunological consequences and underlying mechanisms of early-life wildfire smoke exposure.

Previous studies in rhesus macaques exposed experimentally to ozone and allergens^16–18^, or to natural wildfire smoke^19^, have shown that postnatal exposure during the first three months of life (a critical developmental window for maturation of immune competence, corresponding to the first year of human life^20^) can result in persistent, sex-specific alterations in immune function. Follow-up studies on wildfire-exposed females into adulthood demonstrated that immune abnormalities persisted, albeit with distinct features, and further showed that immune dysregulation extended to their offspring^21^. Together, these findings established that early-life wildfire smoke exposure can have long-lasting immunological consequences. However, these studies were based on the analysis of a limited number of cytokines and genes and did not examine cell type-specific and genome-wide transcriptional and epigenetic mechanisms that contribute to persistent immune dysfunction.

In this study, we leverage a cohort of late-adolescent rhesus macaques at the Oregon National Primate Research Center (ONPRC) that were naturally exposed during infancy to record-breaking hazardous wildfire smoke for nine consecutive days during the 2020 Oregon Labor Day wildfires^22^, to investigate the long-term immune consequences of early-life wildfire smoke exposure. We integrated *ex vivo* immune stimulation, multiplex cytokine quantification, single-cell transcriptomics, and epigenetic profiling, and demonstrate that early-life wildfire smoke exposure is associated with persistent alterations in immune function, immune cell composition, gene regulation, and DNA methylation years after exposure. Together, our findings demonstrate that a single severe wildfire smoke exposure during early life can durably reshape immune cell composition, function, gene regulation, and epigenetic regulation, highlighting acute wildfire smoke exposure as a potential determinant of long-term immune health in the pediatric population.

## RESULTS

### Early-life wildfire smoke exposure has minimal impact on longitudinal growth

Our study included 23 rhesus macaques (Macaca mulatta) housed at the Oregon National Primate Research Center (ONPRC), including 22 of Indian origin and one Indian-Chinese hybrid (Supplemental Table S1). Animals were assigned to either a high wildfire smoke exposure cohort (High-WFS) or a low wildfire smoke exposure cohort (Low-WFS) based on estimated exposure to smoke from the 2020 Oregon Labor Day wildfires (September 9-20, 2020^22^) during early postnatal life (Supplemental Table S1). The High-WFS cohort consisted of animals (n = 6 male, n = 9 female) born within 0-3 months of the 2020 Oregon Labor Day wildfires and housed outdoors during the wildfire event (September 9-20, 2020; Fig. 1A). The Low-WFS cohort consisted of colony counterparts (n = 4 male, n = 4 female) born after the wildfires, as well as one animal born before the fires that was housed in a HEPA-filtered indoor facility during the smoke episode. In the first three months of postnatal life, a critical immune development period in rhesus macaques, High-WFS animals experienced significantly greater cumulative PM_2.5_ exposure, more days with “unhealthy” air quality (PM_2.5_ > 55.4 µg/m³), and higher peak 24hr PM_2.5_ concentrations compared to Low-WFS animals (Table 1). After the first 3 months of life, average and maximum daily PM2.5 exposure were comparable between cohorts (Table 1), highlighting exposure to the wildfire smoke event as the primary environmental exposure distinguishing the two groups. Although High-WFS animals accumulated greater cumulative PM_2.5_ exposure beyond 3 months of age, this was primarily attributable to the modest 2.4-month difference in age between cohorts at time of blood collection (Table 1).

**Figure 1.**
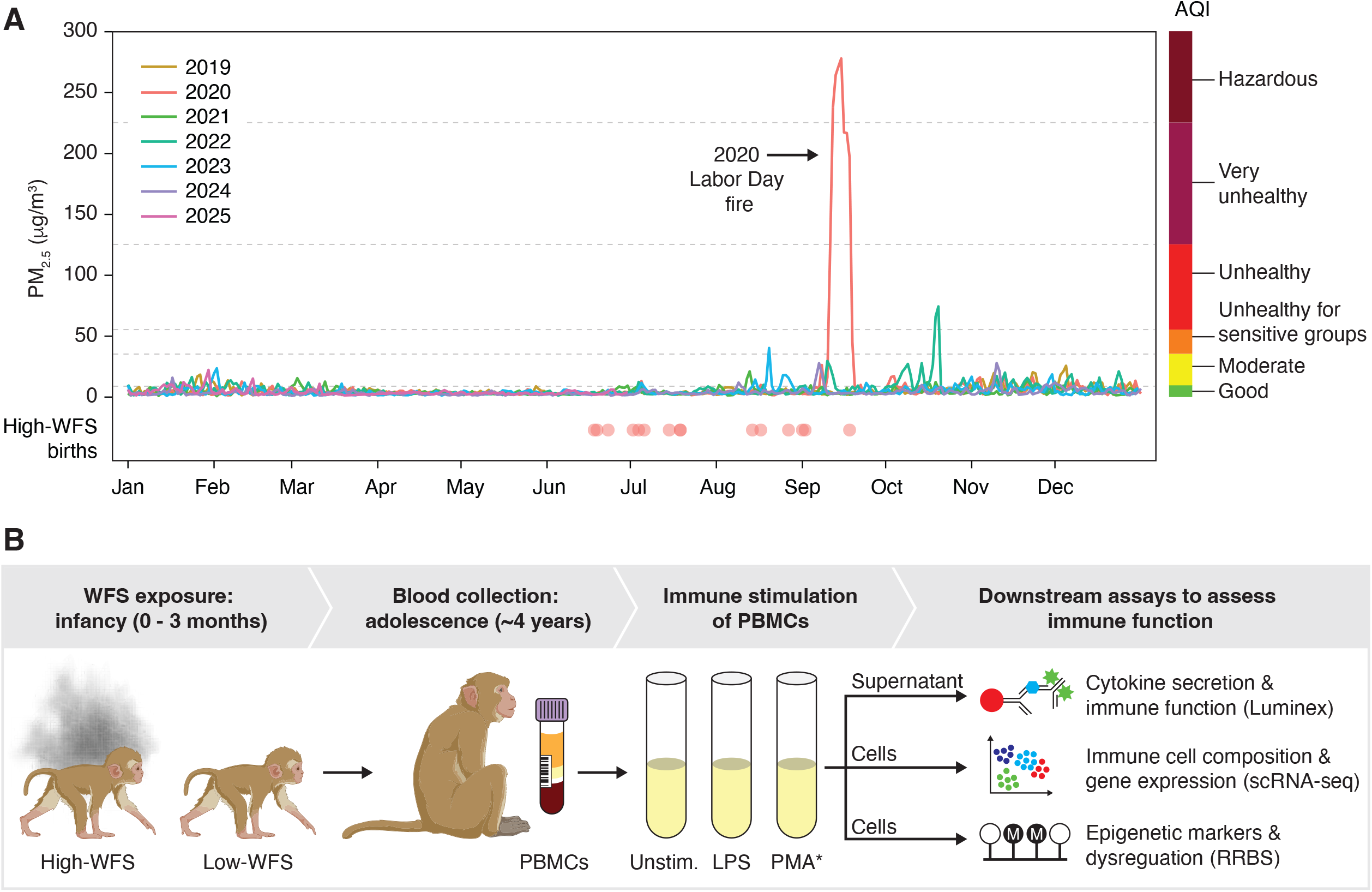
Rhesus macaques exposed to smoke from the 2020 Labor Day wildfire in early postnatal life were used to investigate long-term immune dysregulation. **A)** Daily average PM_2.5_ levels show a marked increase in air pollution near the ONPRC during the 2020 Labor Day wildfires. Birth dates of high-WFS animals are indicated below. Colors on the right correspond to EPA Air Quality Index categories. B) Schematic overview of the study design. Unstim = unstimulated.

**Table 1.**
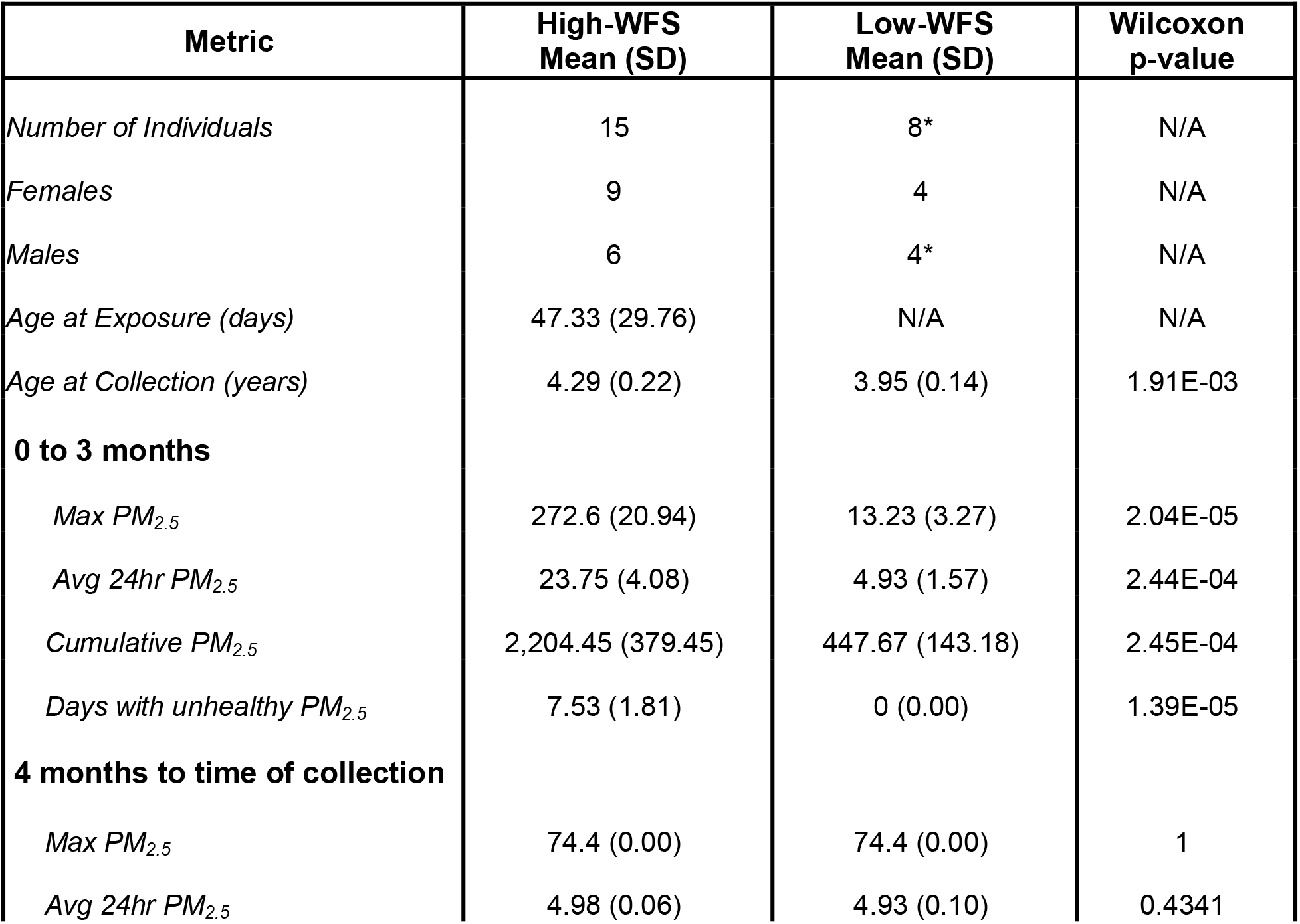
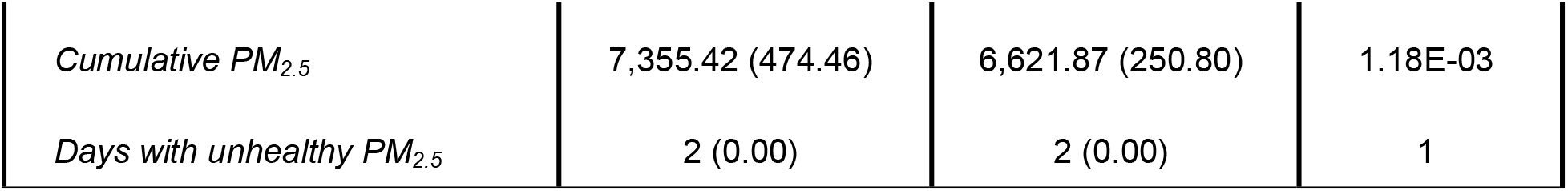
Demographic information for High-WFS and Low-WFS cohorts in this study. *One Low-WFS individual housed in HEPA-filtered facility was excluded from PM_2.5_ statistics reported, since indoor exposure could not be accurately estimated.

Prior studies have reported associations between early-life pollutant exposure and altered birth weight, growth, and metabolic outcomes^23–25^. Therefore, we sought to determine whether wildfire smoke exposure was associated with persistent differences in postnatal growth. We used longitudinal body weight records for all animals, available through veterinary assessment at ONPRC, to investigate long-term weight gain trajectories as a function of early-life wildfire smoke exposure. As expected, weight increased significantly with age (p < 0.0001) in all animals and growth trajectories differed between males and females (age by sex interaction, p < 0.0001). While wildfire smoke exposure was not associated with an overall difference in body weight (p = 0.84) or growth rate (age by WFS exposure interaction, p = 0.96), a significant age by sex by exposure interaction was observed (p = 0.0059), indicating that the association between exposure and growth trajectory differed by sex. Specifically, High-WFS females exhibited slightly slower growth rates compared with Low-WFS females, whereas High-WFS males exhibited the opposite pattern. However, the magnitude of these differences was modest (Supplemental Fig. S1), and overall growth trajectories were broadly similar between exposure groups in both sexes, indicating limited evidence for a biologically meaningful effect of early-life wildfire smoke exposure on longitudinal weight gain in our cohort.

### LPS-induced pro-inflammatory responses are persistently attenuated following early-life wildfire smoke exposure

To investigate the long-term impact of early-life wildfire smoke exposure on the immune system, we collected peripheral blood mononuclear cells (PBMCs) from High- and Low-WFS rhesus macaques ∼4 years after the 2020 Labor Day wildfire when animals were late adolescents (∼4 years of age; Fig. 1B). To evaluate immune function in different contexts, PBMCs from each animal were divided into three experimental conditions: (i) unstimulated, to approximate the resting functional state of circulating immune cells at collection prior to immune challenge (ii) stimulation with lipopolysaccharide (LPS) a Toll-like receptor 4 (TLR4) agonist that models the innate pro-inflammatory response to bacterial infection, and (iii) stimulation with phorbol 12-myristate 13-acetate (PMA; without ionomycin) to assess early activation responses downstream of protein kinase C (PKC) signaling (Fig. 1B).

As a broad assessment of immune function, we quantified secretion of 27 cytokines in culture supernatants following 10-hours *ex vivo* stimulations. As expected, LPS exposure induced robust proinflammatory cytokine responses in both sexes, with significant increases in IL-6, IL-8, IL-2, CCL4, CD40 ligand, GM-CSF, IFN-α, IL-1β, PD-L1, PDGF, and TNF-α (Wilcoxon signed-rank test p < 0.05; Fig. 2A; Supplemental Table S2). Only in females LPS also induced significant changes in Granzyme B, IL-10 and CCL5 (Wilcoxon signed-rank test p<0.05), while males exhibited similar but non-significant trends for these cytokines. In contrast to LPS, PMA elicited modest and less consistent responses, with significant changes detected for Granzyme B in both sexes, for IL-8 and PDGF-BB in females, and for Granzyme B in males (Wilcoxon signed-rank test p <0.05; Supplemental Table S2).

**Figure 2.**
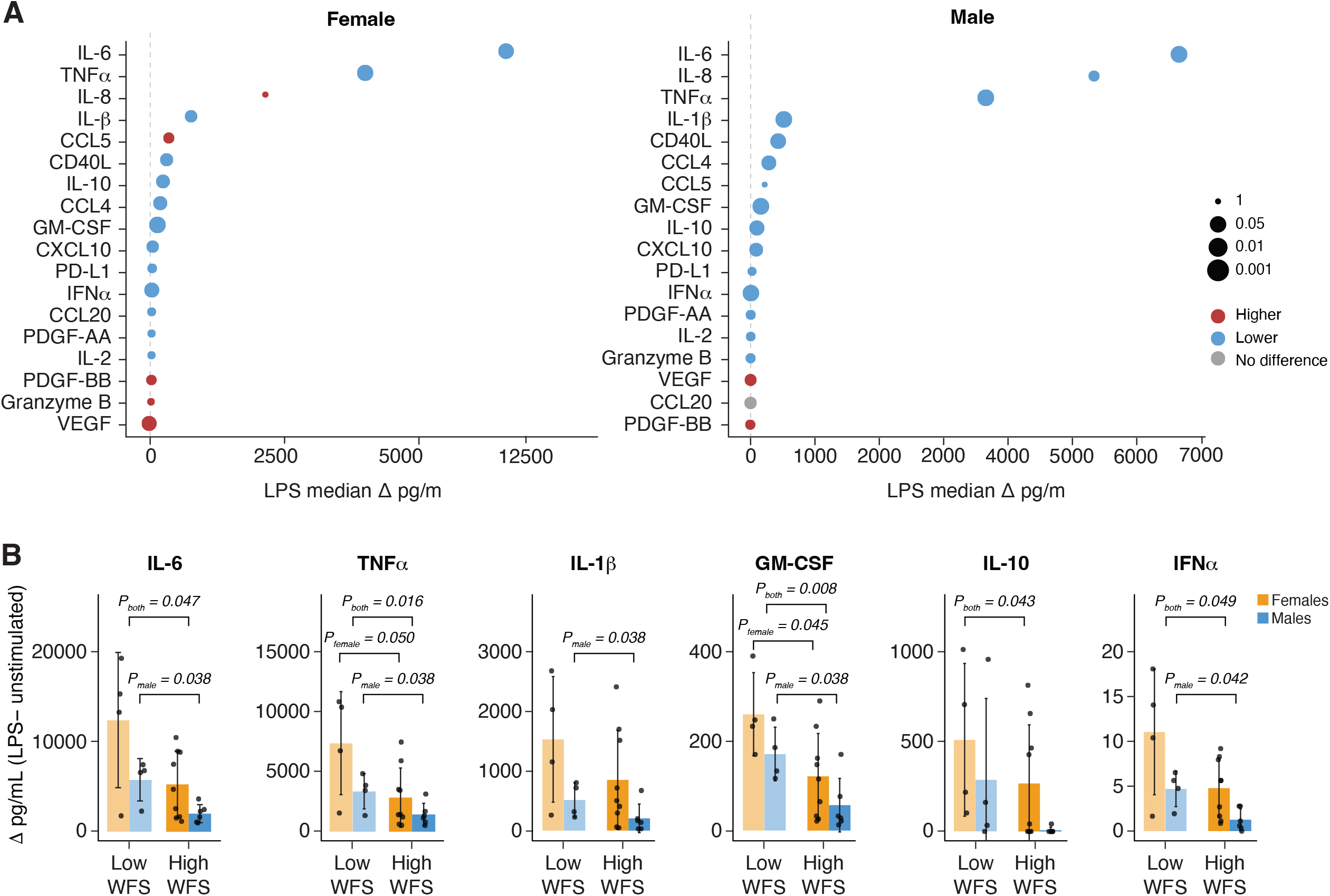
Early postnatal wildfire smoke exposure is associated with attenuated pro-inflammatory cytokine responses. **A)** Forest plots show the overall direction and magnitude of background-corrected cytokine response to LPS stimulation for Low-WFS females and males. Color represents relative direction of changes observed in High-WFS animals. **B)** Background-corrected quantification of cytokines significantly altered in high-WFS animals following LPS stimulation. Only p-values ≤0.05 are shown (mean+/-stdev).

To determine whether early-life wildfire smoke exposure altered immune responsiveness, we compared background-corrected cytokine responses (Δ = stimulated − unstimulated) between High-WFS and Low-WFS animals using both sex-stratified and sex-combined analyses. Among cytokines responsive to LPS stimulation, IL-6, TNF-α GM-CSF, IL-10, and IFN-α showed significantly reduced secretion in High-WFS animals in sex-combined analyses (Wilcoxon signed-rank test p < 0.05; Supplemental Table S2). Sex-stratified analyses suggested a stronger attenuation of the LPS response among High-WFS males, with significant reductions in IL-1β, TNF-α, IFN-α, IL-6, GM-CSF (Fig. 2B). Although all five cytokines showed the same direction of change in High-WFS females, only GM-CSF reached statistical significance. In contrast to LPS, PMA stimulation was associated with limited exposure-associated differences, with only VEGF displaying significant reduction in High-WFS animals in sex-combined analysis (Wilcoxon signed-rank test p = 0.03; Supplemental Table S2). Together, these findings indicate persistent attenuation of LPS-induced innate inflammatory responses following early-life wildfire smoke exposure, with the strongest effects observed in males.

### Early-life wildfire smoke exposure is associated with persistent alterations in composition and transcriptional programs across immune cell types

Given the persistent differences in innate immune response associated with early-life wildfire smoke exposure, we next used single-cell RNA sequencing (scRNA-seq) to investigate long-term alterations in peripheral immune cell composition and cell-specific gene expression patterns. PBMCs collected from the 23 rhesus macaques under unstimulated and LPS-, or PMA-stimulated conditions produced 69 scRNA-seq libraries totaling 185,910 cells. Following quality control filtering, doublet removal, and exclusion of non-immune, dead or low-quality cells, 129,467 cells were retained for downstream analysis. Using annotations based on the Rhesus Immune Reference Atlas or RIRA^26^ cells were classified into three populations: myeloid cells (n=12,719), B cells (n=39,416), and T/Natural Killer (NK) cells (n=73,470; Fig. 3A). Cells classified as “Unknown” (n = 3,862) were excluded from subsequent analyses. The myeloid group encompasses cells of the myeloid lineage and is expected to be composed primarily of monocytes and dendritic cells. T cells and NK cells, although distinct immune cell populations, share many effector functions and exhibit similar transcriptional profiles. Consequently, they are grouped into a common T/NK compartment^26^ (Fig. 3A). Comparison of immune cell composition revealed significant shifts in High-WFS males, associated with increased proportions of T/NK cells and reduced B cells (Fig. 3B). These differences were evident at unstimulated state, as well as following both LPS and PMA stimulation (Supplemental Fig. S2A).

**Figure 3.**
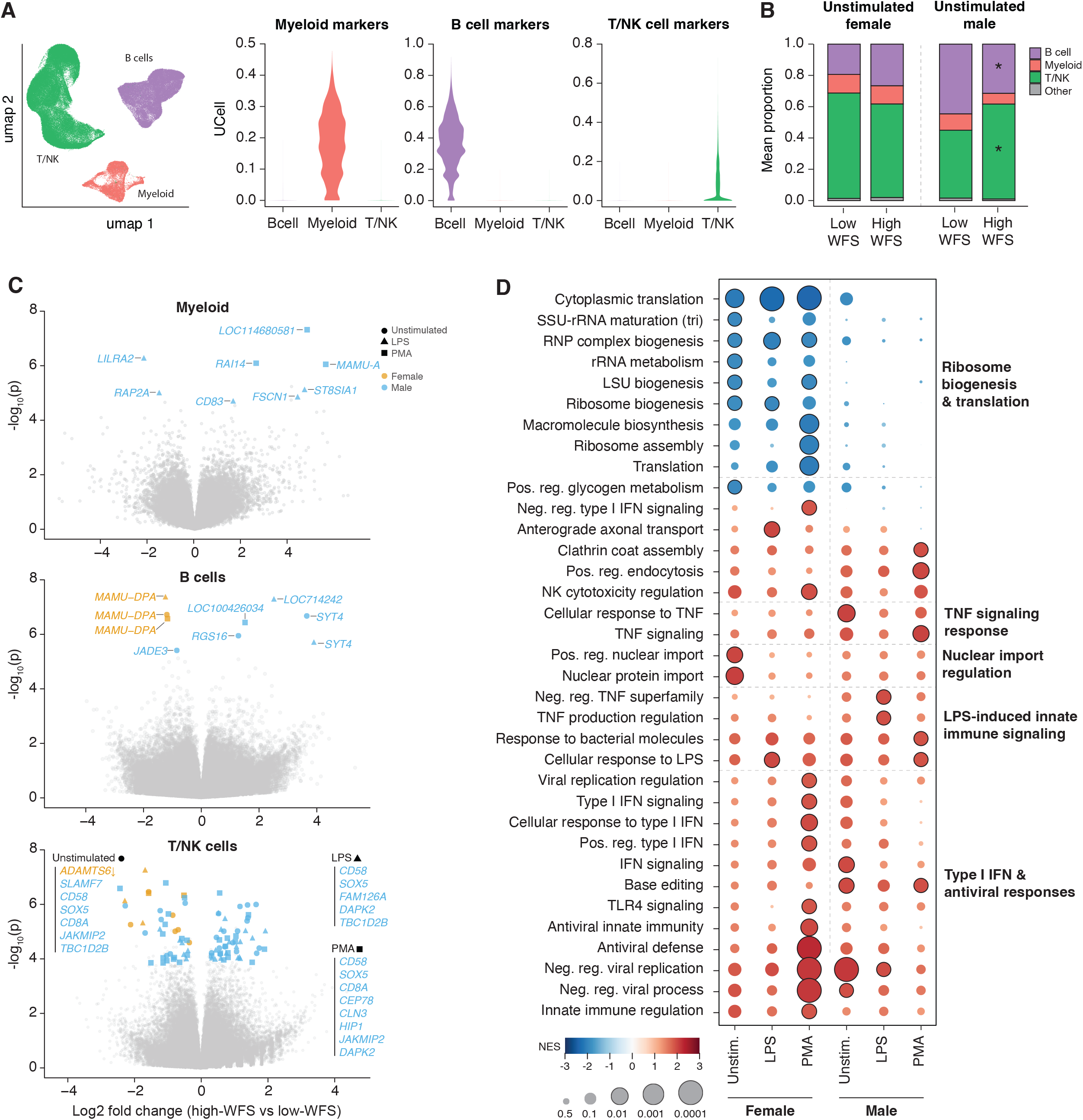
Long-term gene dysregulation is detected across major immune cell types of High-WFS animals. **A)** Lineage-specific gene expression signatures are used by RIRA to annotate B cell, Myeloid and T cell. **B)** High-WFS males exhibit increased T/NK and decreased B cell proportions relative to Low-WFS. **C)** Volcano plots show DEGs (FDR q ≤ 0.05) between High- and Low-WFS animals within each cell type (male DEGs in blue and female in orange). Symbol shape indicates the condition (unstimulated, LPS, or PMA) in which the DEG was identified. For T/NK cells, only DEGs with ≥%20 of T/NK cells expressing the gene and |log2FC| ≥ 0.5 are labeled. **D)** GSEA analysis reveals coordinated pathway dysregulation in T/NK cells in males and females. Related pathways are grouped based on gene overlap. Color indicates GSEA Normalized Enrichment Score (NES) in High-WFS relative to controls. Dot size reflects statistical significance, with those with q<0.05 outlined in black. Unstim = unstimulated.

To determine whether the attenuated inflammatory response based on cytokine profiling was also reflected at the transcriptional level, we generated cell type-specific inflammatory and TNFα-response signatures from genes significantly induced by LPS or PMA that overlapped with the MSigDB Hallmark^27^ “Inflammatory Response” and “TNFα Signaling via NF-κB” gene sets (Supplemental Table S3). We found subtle sex-specific attenuation of inflammatory transcriptional responses in myeloid cells from High-WFS males following LPS stimulation. Collectively, High-WFS male myeloid cells exhibited significantly reduced median enrichment of LPS-responsive TNFα pathway genes, compared to Low-WFS males (p = 0.026, Student’s t-test), with a similar trend observed for the 75th percentile of enrichment (p = 0.078, Student’s t-test; Supplemental Table S3). Likewise, LPS-responsive inflammatory pathway genes showed reduced enrichment in High-WFS males, including a trend toward lower median UCell (i.e., rank-based gene signature enrichment^28^) scores (p = 0.13, Student’s t-test) and a significant reduction in the 75th percentile UCell score (p = 0.03, Student’s t-test; Supplemental Table S3). No differences were observed in any setting following PMA stimulation, in female subjects, or within T/NK or B cell populations for either pathway’s gene set (Supplemental Table S3). Together, these results suggest that early-life wildfire smoke exposure is associated with a persistent reduction in the magnitude of myeloid inflammatory responses to LPS, rather than a reduction in the number of responsive cells.

We next performed differential expression analysis on pseudobulk RNA-seq data from each coarse immune cell type, to identify transcriptional alterations associated with early-life wildfire smoke exposure. Males and females were analyzed separately motivated by previously reported sex differences in smoke-induced immune dysregulation in rhesus macaques^19^, as well as the sex differences observed in our own cytokine data. Within myeloid cells we identified a total of eight differentially expressed genes (DEGs) in males only (adjusted p ≤ 0.05; Fig 3C; Supplemental Table S4). While limited in number, these DEGs highlighted biologically relevant pathways involved in innate immune activation, TLR inflammatory signaling, and antigen presentation. Under LPS stimulation, myeloid cells of High-WFS males exhibited significantly reduced expression of *LILRA2* (log2FC = −2.18, padj = 0.007), an immunoglobulin-like receptor involved in myeloid cell activation and TLR-mediated inflammatory^29^, and *RAP2A* (log2FC = −1.52, padj = 0.04), a small Ras-related GTPase implicated in TLR- and LPS-mediated NF-κB signaling^30^. In contrast, High-WFS males had increased expression of *CD83* (log2FC = 1.6, padj = 0.05), a canonical dendritic cell activation marker also expressed by LPS-activated monocytes and macrophages^31^, and *FSCN1* (log2FC = 4.4, padj = 0.046), which encodes an actin-bundling protein associated with dendritic cell maturation and migration^32^ (Supplemental Table S4). Under PMA stimulation, High-WFS male myeloid cells exhibited upregulation of *MAMU-A* (log2FC = 5.62, padj = 0.004), a rhesus macaque MHC class I gene involved in antigen presentation to CD8^+^ T cells, suggesting altered antigen-presentation activity. No significant DEGs (padj ≤ 0.05) were detected in unstimulated conditions in male myeloid cells or in females under any condition (Fig. 3C; Supplemental Table 4). Collectively, these transcriptional changes are consistent with the attenuated LPS-driven inflammatory cytokine responses observed in High-WFS males, while suggesting that wildfire smoke exposure may differentially affect inflammatory signaling and other activation-associated aspects of the myeloid response to LPS, such as dendritic cell maturation.

Similar to myeloid cells, relatively few DEGs were identified in B cells following early-life wildfire smoke exposure, with only five unique DEGs detected across the three stimulation conditions and two sexes (Fig. 3C; Supplemental Table S4). Of note, *MAMU-DPA*, the rhesus macaque ortholog of human *HLA-DPA1* encoding an MHC class II antigen-presentation molecule, was downregulated in female B cells across all three conditions (log2FC = −1.2, padj < 0.01). In males, *RGS16,* a regulator of chemokine signaling and lymphocyte trafficking^33^, was upregulated in unstimulated condition (log2FC = 1.27, padj = 0.02; Fig. 3C; Supplemental Table S4). These findings suggest that early-life wildfire smoke exposure might also impact B cell antigen presentation and immune signaling pathways in a sex-specific manner.

In contrast to B cells and myeloid cells, T/NK cells exhibited more extensive transcriptional alterations associated with early-life wildfire smoke exposure (Supplemental Table S4). Female High-WFS animals displayed fewer changes, with eight unique DEGs identified across all conditions, all of which were downregulated, including, *HMOX2*, which encodes heme oxygenase 2, a key regulator of heme metabolism and cellular responses to oxidative stress^34^, was reduced following PMA stimulation (log2FC = −0.52, Fig. 3C; adjusted padj = 0.007). In addition, *UBAC2*, a ubiquitin-associated protein implicated in immune regulation and autoimmune disease susceptibility^35^, was decreased under the unstimulated condition (log2FC = −0.36, padj = 0.05). Among males, the High-WFS cohort exhibited a broader transcriptional change, with 52 unique DEGs identified across all conditions (Fig. 3C; Supplemental Table S4). Among the most notable was *SOX5*, a transcription factor involved in T cell differentiation and signaling^36^, which was consistently upregulated across all three conditions (log2FC = 1.17-1.40, padj < 0.05). *SLAMF7*, encoding an immunoregulatory surface receptor found on NK cells and some T cells, and implicated in activation and effector function^37^, was significantly increased at unstimulated condition (log2FC = 1.06, padj = 0.04). We also observed elevated expression of *CD58*, a co-stimulatory adhesion molecule that promotes T cell activation through interaction with CD2^38^, across all three stimulation conditions (log2FC = 0.41-0.55, padj < 0.05). Notably, *CD8A*, which encodes the alpha chain of the CD8 co-receptor and serves as a marker of cytotoxic T lymphocytes, was significantly increased at unstimulated and PMA stimulated conditions (log2FC ≈ 0.7, padj = 0.03) with a similar trend following LPS stimulation (log2FC ≈ 0.6, padj = 0.07; Fig. 3C). Although the specific DEGs reaching statistical significance differed between males and females, the overall direction of expression changes was broadly concordant across sexes, with male animals generally exhibiting larger effect sizes and stronger statistical support (Supplemental Fig. S2B).

Gene set enrichment analysis (GSEA^39^) of DEG in the T/NK compartment revealed broad pathway-level alterations associated with wildfire smoke exposure (Supplemental Table S5). While the specific pathways reaching statistical significance differed between sexes, the overall direction of enrichment was remarkably consistent (Fig. 3D; Supplemental Table S5). High-WFS animals showed reduced expression of several pathways involved in cellular biosynthetic activity, while several other pathways related to innate immune signaling, responses to bacterial products and LPS, type I interferon and antiviral responses and regulation of NK-cell-mediated cytotoxicity were increased (Fig. 3D). Collectively, these findings indicate long-lasting transcriptional reprogramming of T/NK cells following early-life wildfire smoke exposure in both sexes.

### Early-life wildfire smoke exposure is associated with higher proportion of cytotoxic T/NK cells and transcriptional alterations in CD8^+^ cells

Motivated by the differential expression of genes (e.g. *CD8A*) and pathways implicated in T/NK cell effector function and cytotoxicity in High-WFS animals, we asked whether cytotoxic T/NK cells were specifically impacted by early-life wildfire smoke exposure. We observed a significant increase in the fraction of T/NK cells expressing cytotoxic gene program (UCell score > 0) in High-WFS animals at across all three stimulation conditions, in both sexes (Fig. 4A). Among T/NK cells expressing the cytotoxic program, median cytotoxicity scores also trend higher in High-WFS animals at unstimulated and PMA stimulated conditions, indicating modest context-dependent increases in the intensity of cytotoxic gene expression (Fig. 4B).

**Figure 4.**
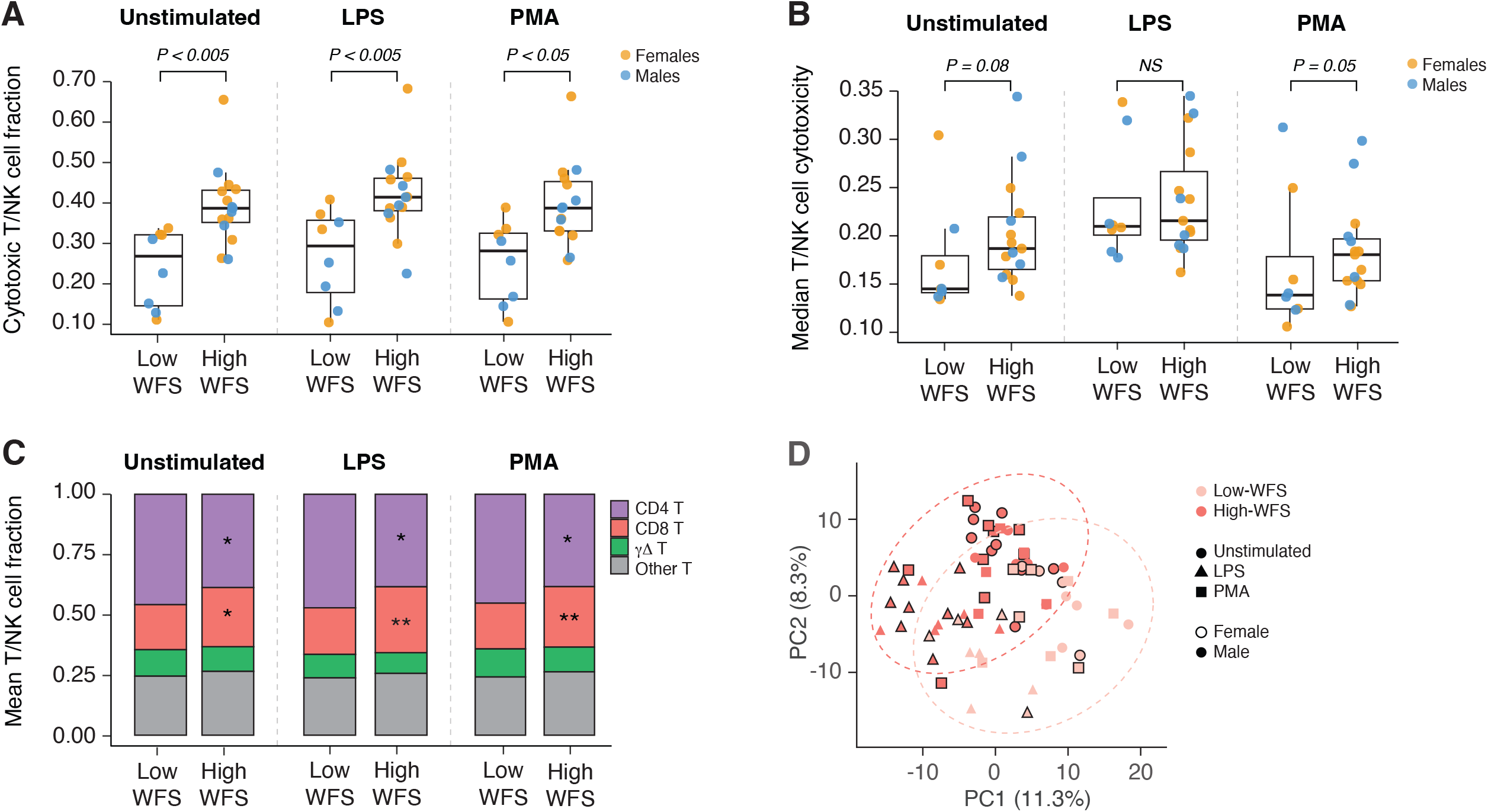
Cytotoxic T cells are expanded and transcriptionally altered following early postnatal wildfire smoke exposure. **A)** A greater fraction of T/NK cells exhibit cytotoxic gene program enrichment (UCell > 0) in male and female High-WFS macaques. **B)** Median cytotoxicity among UCell-positive cells is modestly increased in High-WFS animals. **C)** RIRA annotations reveal a higher proportion of CD8+ T cells and a lower proportion of CD4^+^ T cells in High-WFS animals across conditions (*p < 0.05, **p < 0.005). **D)** PCA of the 500 most variable genes in CD8^+^ T cells separates subjects by early postnatal wildfire smoke exposure status.

To further characterize T cell subpopulations, we used RIRA^26^ to classify T cells into CD4^+^, CD8^+^ and γδ cells. Unclassified T-cells were excluded from downstream analyses because they lacked confident annotation, and γδ T cells were excluded because of their low abundance (Fig. 4C). Consistent with the increased abundance of cytotoxic T/NK cells, differential cluster abundance analysis identified a significant increase in the proportion of CD8^+^ T cells in High-WFS animals (Fig. 4C). We next examined whether wildfire smoke exposure was associated with transcriptional alterations within the CD8^+^ and CD4^+^ compartments. UMAP visualization of CD8^+^ T cells based on the 500 most variable genes revealed separation by smoke exposure status (Fig. 4D). Consistent with this, differential expression analysis identified widespread transcriptional changes in CD8^+^ T cells of High-WFS animals. In males, 101 unique DEGs (padj ≤ 0.05) across conditions were identified and in females we identified 367 unique DEGs across conditions. Of these DEGs, 24 were shared between males and females, included several genes with established roles in T cell activation and differentiation; For example, *AIM2*, an innate immune sensor involved in inflammatory responses^40^, *BACH2,* a transcription factor that maintains T cell in naïve states^41^, *ZEB1*, a regulator of T cell differentiation and memory formation^42^, and *PTPN22,* a key regulator of T cell receptor signaling^43^ (Fig. 5A; Supplemental Table S4). Most genes were identified exclusively in one sex, for example female High-WFS animals exhibited reduced expression of *FOXP1* and decreased expression of *THEMIS*, both genes implicated in T cell development and signaling^44,45^ (Fig. 5A). Male High-WFS animals showed increased expression of *ZEB2* and elevated *RUNX1*, genes associated with lymphocyte differentiation and effector cell development^42,46^ (Fig. 5A; Supplemental Table S4).

**Figure 5.**
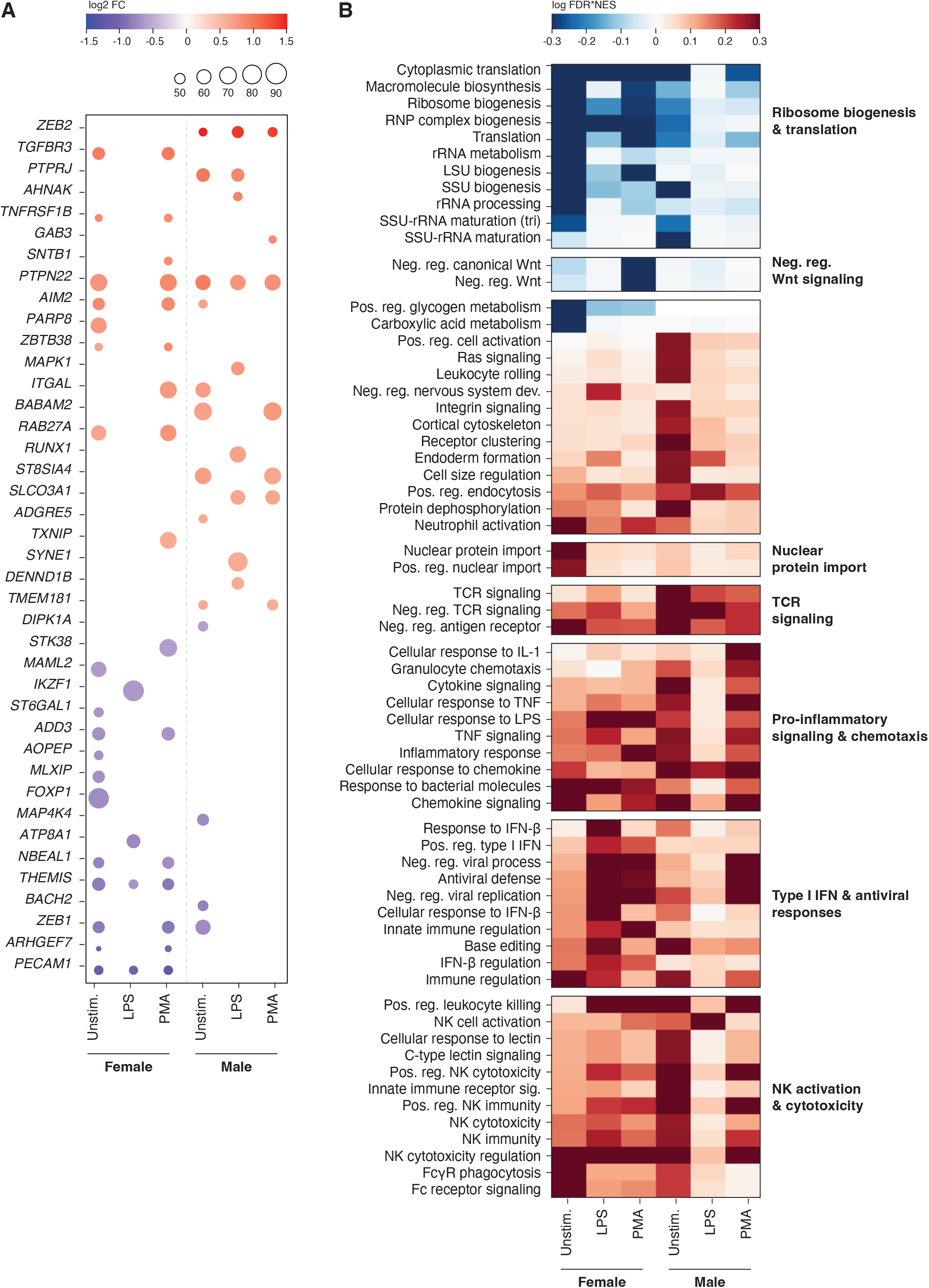
Genes and pathways are persistently dysregulated in CD8^+^ T cells of High-WFS animals. **A)** Numerous genes are differentially expressed in CD8^+^ T cells from WFS animals. Dot color indicates log2FC (capped at ±1.5), and dot size indicates the percentage of CD8^+^ T cells expressing each gene. Only genes with |log2FC| ≥ 0.6, FDR < 0.06, and expression in ≥45% of CD8^+^ T cells are shown. **B)** GSEA reveals coordinated pathway-level dysregulation in CD8+ T cells. Related pathways were grouped based on gene overlap. Red indicates higher and blue indicates lower expression in High-WFS animals. Unstim = unstimulated.

Gene set enrichment analysis of DEG revealed broad pathway-level alterations that were largely consistent between males and females despite differences in individual DEGs. High-WFS animals exhibited reduced expression of pathways related to ribosome biogenesis, protein translation, and WNT signaling. In contrast, pathways associated with pro-inflammatory signaling, chemotaxis, type I interferon and antiviral responses, and NK-cell activation and cytotoxicity were enriched in smoke exposed animals (Fig. 5B; Supplemental Table S5). In contrast to the extensive transcriptional changes observed in CD8^+^ T cells, no exposure-associated differentially expressed genes were detected within CD4^+^ T cells (Supplemental Fig. S4), indicating that cytotoxic T/NK cells are particularly susceptible to long-term dysregulation following early postnatal wildfire smoke exposure.

### Early-life wildfire smoke exposure is associated with increased CD8^+^ T cell differentiation, with the greatest effects following exposure at younger ages

Considering that naïve-to-memory differentiation is a major source of transcriptional variation in T cells, and that several DEGs we identified in CD8^+^ T cells are implicated in differentiation and effector function, we investigated differentiation levels of CD4^+^ and CD8^+^ T cells using the Effector Differentiation Score (EDS), a validated gene expression-based metric of peripheral T cell differentiation^26^. As expected, EDS values displayed a gradient across T cells, consistent with the presence of cells spanning a spectrum of differentiation (Fig. 6A). Inspection of overall cell EDS distributions revealed a clear collective shift toward higher differentiation scores in CD8^+^ T cells in both male and female High-WFS animals relative to Low-WFS controls (Fig. 6B). Using previously established EDS thresholds^26^, we classified CD8^+^ and CD4^+^ T cells as naïve (T_n_ EDS < 2), central memory (2 ≤ T_cm_ EDS < 6), or effector memory (T_em_ EDS ≥ 6; Fig 6B). Across unstimulated, LPS, and PMA conditions, High-WFS animals exhibited a significant reduction in the proportion of naïve CD8^+^ T cells accompanied by a corresponding increase in effector memory CD8^+^ T cells, while the frequency of central memory cells remained unchanged (Fig. 6C). These findings indicate a persistent shift of the CD8^+^ compartment, regardless of sex, toward a more differentiated phenotype following early-life wildfire smoke exposure. In contrast, no differences in differentiation were observed among CD4^+^ T cells (Supplemental Fig. S3).

**Figure 6.**
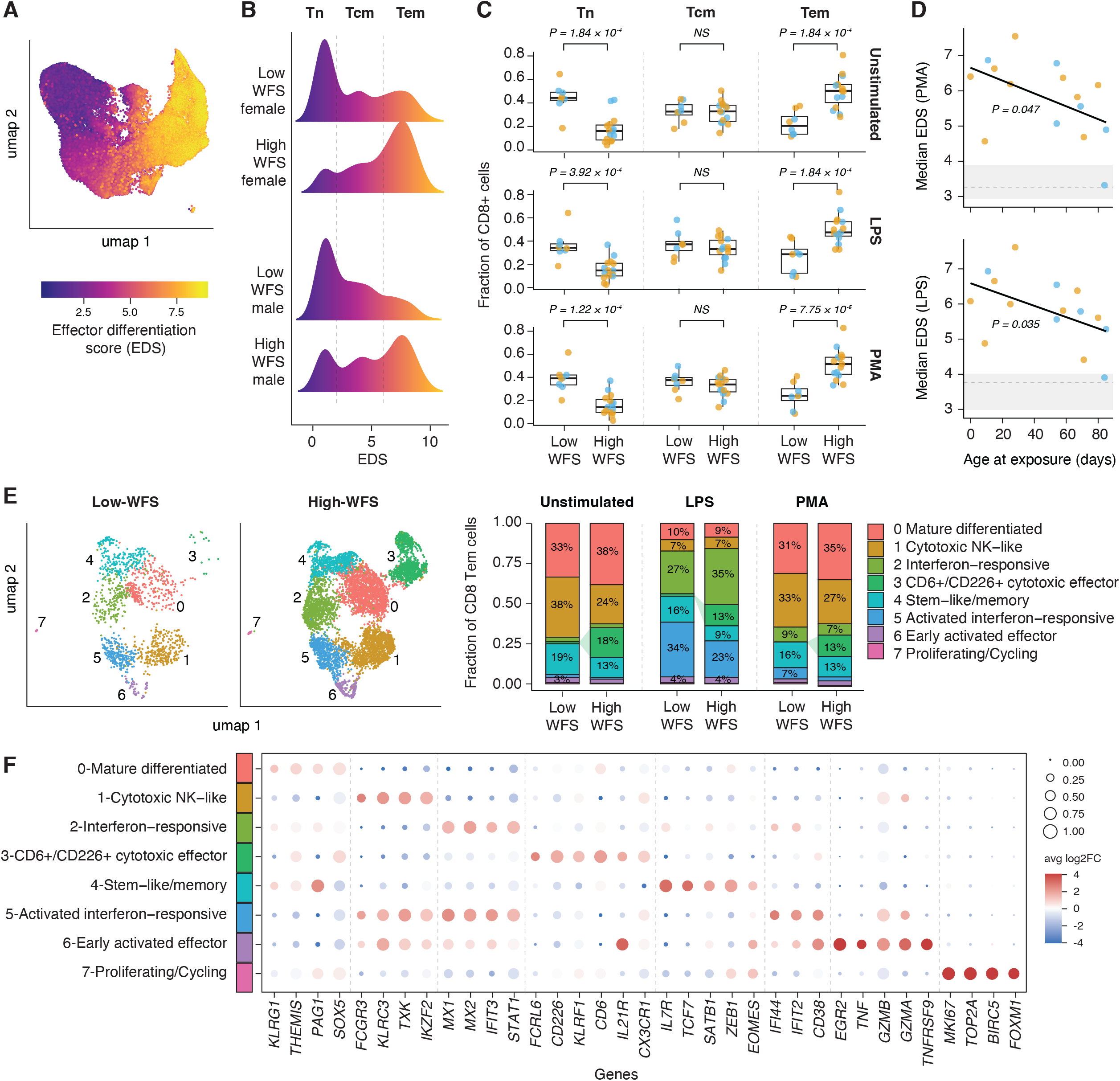
CD8^+^ T cells in High-WFS display a shift toward more differentiation and expansion of a distinct Tem population. **A)** The Effector Differentiation Score (EDS) quantifies T cell differentiation along a continuous trajectory from naïve to effector states. **B)** Density distributions of CD8^+^ cells along the EDS trajectory reveal a shift toward more differentiated effector states in both male and female High-WFS animals. CD8^+^ cells were categorized as naïve (T_n_), central memory (T_cm_), and effector memory (T_em_) based on EDS value. **C)** High-WFS animals exhibit a significantly lower fraction of T_n_ cells and a higher fraction of T_em_ cells across all conditions. **D)** Under LPS and PMA stimulation, median EDS is negatively correlated with age at wilddfire smoke exposure among High-WFS animals. The interquartile range and median of Low-WFS animals are shown in gray. **E)** Left, UMAP visualization of T_em_ CD8^+^ cells from High- and Low-WFS animals identifies eight transcriptionally distinct clusters. Right, average cluster composition in High- and Low-WFS animals is shown. **F)** Select marker genes are displayed for each cluster. Color indicates average expression enrichment (log2FC), and dot size represents the percentage of cells expressing the gene within each cluster.

Notably, the degree of CD8^+^ T-cell differentiation in High-WFS animals under LPS and PMA stimulation, quantified by the median EDS, was significantly associated with age at wildfire smoke exposure (unstimulated: Spearman’s ρ = −0.36, p = 0.18; LPS: Spearman’s ρ = −0.54, p = 0.036; PMA: Spearman’s ρ = −0.52, p = 0.048; Fig. 6D). In other words, High-WFS animals exposed to wildfire smoke at younger ages exhibited a greater shift toward differentiated CD8^+^ T cell states. Because animals age at smoke exposure was broadly correlated with age at the time of blood collection (Spearman’s ρ = 0.66, p = 0.0078), we evaluated whether age at sampling rather than age at exposure could explain the observed association with CD8^+^ T-cell differentiation. We found that regression models incorporating either age at exposure or age at blood collection showed similar fit (AIC: 134.1 vs. 134.9, respectively). However, among Low-WFS animals, age at blood collection was not associated with median CD8^+^ EDS, despite spanning a similar age range (Spearman p > 0.05; Supplemental Fig. S3). Together, these findings suggest that the association with CD8^+^ T-cell differentiation is more consistent with an effect of the timing of wildfire smoke exposure.

To determine whether the increased abundance of CD8^+^ effector memory (T_em_) cells in High-WFS animals reflected expansion of a novel differentiated cell state or a broad shift toward effector differentiation, we subset all CD8^+^ T cells with EDS > 6, a threshold previously shown to capture effector memory T cells^26^. PCA/UMAP analysis of this subset identified eight transcriptionally distinct clusters (Fig. 6E). Under unstimulated (and PMA-stimulated) conditions, High-WFS animals exhibited modest compositional shifts consistent with increased differentiation, including a reduced proportion of stem-like/memory cells (13% vs. 19% in cluster 4) and an increased proportion of differentiated effector cells (38% vs. 33% in cluster 0; Fig. 7E). In addition, we identified a distinct cluster (cluster 3) of cells exclusively present among High-WFS animals and characterized by elevated expression of *FCRL6*^47^, *CD226*^48^, *KLRF1*^49^, *CD6*^50^, *IL21R*^51^, and *CX3CR1*^52^, genes associated with cytotoxic lymphocyte function, activation, and effector differentiation. We termed this population the CD6^+^/CD226^+^ cytotoxic effector cluster (Fig. 6F). This population was nearly absent in Low-WFS animals but comprised 13-18% of T_em_ cells in High-WFS animals (Fig. 6E), with most of these cells originating from four individuals (three females and one male). Following LPS stimulation, High-WFS showed modest increase in proportion of cells within the interferon-responsive cluster (35% vs. 27% in cluster 2), whereas Low-WFS animals appeared to have a larger fraction of cells within the activated interferon-responsive cluster (34% vs. 23% in cluster 5). Together, these findings suggest that the increased abundance of CD8^+^ T_em_ cells in High-WFS animals likely reflects both a general shift toward more differentiated effector states, as well as expansion of a distinct cytotoxic effector-memory cell population.

**Figure 7.**
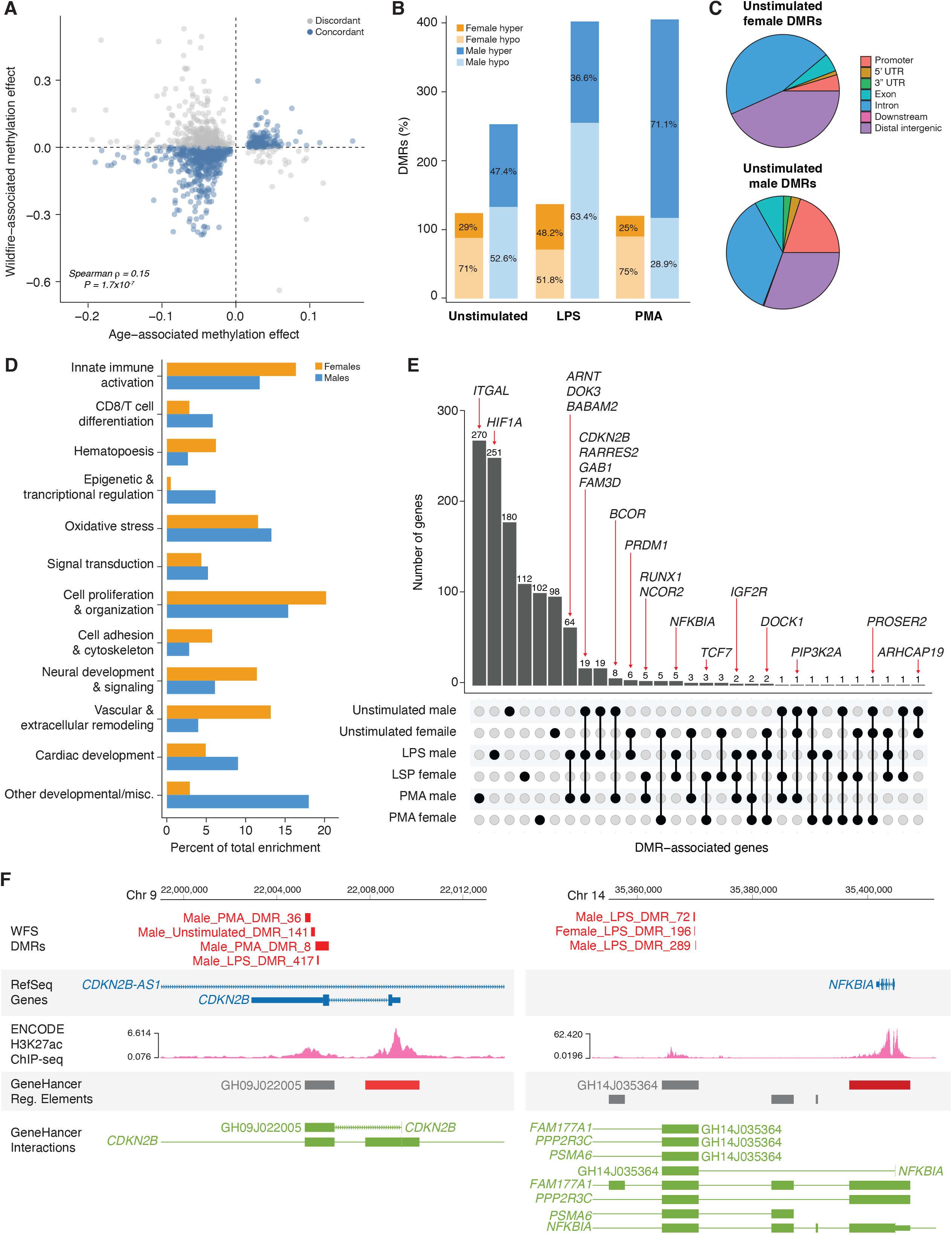
Early-life wildfire smoke exposure induces persistent epigenetic changes in immune genes and regulatory elements. **A)** Age-associated CpG sites exhibiting concordant versus discordant methylation changes following wildfire smoke exposure, with a bias toward concordant changes consistent with accelerated epigenetic aging. **B)** Number of DMRs and the proportion of hypo- and hypermethylated regions in each condition. **C)** Distribution of DMRs across genomic features, showing that most occur in noncoding regions. **D)** Percentage of significantly enriched GO pathways grouped by broad biological function. **E)** Upset plot showing overlap of DMR-containing genes across conditions, with selected biologically relevant genes highlighted. **F)** Two hypermethylated loci in High-WFS animals, mapped to hg38, contain DMRs overlapping predicted enhancer elements that interact with their target genes. Tracks show (top to bottom) DMRs, RefSeq genes, ENCODE heart H3K27ac ChIP-seq signal, GeneHancer regulatory elements (enhancers shown in gray), and GeneHancer regulatory interactions.

### Early postnatal wildfire smoke exposure shifts DNA methylation patterns toward an older epigenetic age

Early-life exposure to ambient toxicants, including tobacco smoke and air pollution, have been associated with accelerated epigenetic aging^53,54^. Thus, we asked whether acute early postnatal wildfire smoke exposure was associated with changes in epigenetic aging in our subjects. We used genome-wide DNA methylation profiling of PBMCs from High- and Low-WFS animals across three immune stimulations to estimate epigenetic age based on a recently developed rhesus macaque epigenetic clock^55^. The clock generated accurate age estimates at unstimulated conditions, with predicted ages clustering within the expected range for this cohort and low median absolute error (MAE; Low-WFS = chronological age +/-1.38 years, High-WFS = age +/- 0.78 years). Within this restricted age range, prediction errors of only a few years are consistent with satisfactory clock performance. Across all stimulation conditions, comparison of epigenetic age acceleration (predicted age - expected age) revealed a subtle trend toward older epigenetic age in High-WFS animals relative to Low-WFS (+0.80 years, p = 0.09; Supplemental Table S6). Notably, the magnitude of the difference was greater following immune stimulations than in the unstimulated state, suggesting that inflammatory challenges may accentuate signatures of epigenetic aging.

Because epigenetic clocks compress information from hundreds of loci into a single age estimate and have limited resolution among individuals of similar age, we next examined methylation patterns at independently defined age-associated loci^55^. Under unstimulated condition, 801 of 1,211 (66.1%) of age-associated genomic regions exhibited exposure-associated methylation changes in the same direction as the established age effect. Thus, High-WFS animals displayed methylation patterns that more closely resembled those of older individuals. This directional concordance was significantly greater than the 50% expected by chance (binomial test, p = 0.0001; 95% CI: 63.4-68.8%; Fig. 7A; Supplemental Fig. S4B; Supplemental Table S6). This effect appeared more pronounced at loci that become hypermethylated with age (84.1% directional concordance), compared with loci that undergo hypomethylation (62.4% concordance; χ² = 36.1, p = 1.8 × 10^-9^). We also found that the magnitude of age-associated and exposure-associated methylation effects show a significant positive correlation across loci (Spearman’s ρ = 0.15, p = 1.7 × 10^-7^; Fig.7A). Together, these results provide evidence that early-life wildfire smoke exposure may accelerate epigenetic aging resulting in persistent, genome-wide shifts in DNA methylation toward an older state.

### Persistent DNA methylation changes in High-WFS may contribute to immune dysregulation

The long-term effects of early-life environmental exposures are often mediated by epigenetic mechanisms, including DNA methylation, that can alter gene regulation without modifying the underlying DNA sequence^56,57^. To investigate the potential contribution of DNA methylation to immune dysregulation in our cohort, we leveraged our DNA methylation data to identify all differentially methylated regions (DMRs) between High- and Low-WFS cohorts. Each DMR was annotated to the overlapping gene or, when no overlap existed, to the nearest gene as the most likely candidate target (Supplemental Table S7). Considering DMRs that showed at least 10% difference in DNA methylation between High- and Low-WFS, in males we identified 235, 402, and 405 DMRs under unstimulated, LPS, and PMA conditions, respectively, whereas females exhibited fewer DMRs (124, 137, and 120, respectively; Fig. 7B). Except for male PMA samples, the majority of DMRs in each sex and stimulation condition (52-75%) were hypomethylated in High-WFS animals (Fig. 7B). In both sexes, >90% of DMRs were located within non-coding regions, with nearly half of these DMRs mapping to distal intergenic regions (Fig. 7C). Within distal intergenic DMRs we detected significant enrichment of 12 transcription factor binding motifs (Benjamini-Hochberg padj < 0.05), including motifs recognized by immune regulatory transcription factors such as NR2F6, REV-ERBα, and USF2 (Supplemental Table S8), suggesting that wildfire smoke-associated DNA methylation changes preferentially occur within immune regulatory elements. Consistent with this, liftOver of DMRs to the human genome (hg38) showed that 239 of 1,161 successfully mapped DMRs (20.6%) overlapped at least one candidate cis-regulatory element (cCRE) defined by CATlas single-cell ATAC-seq datasets spanning nine adult human immune cell types (macrophages, alveolar macrophages, CD4^+^ T cells, CD8^+^ T cells, naïve T cells, NK cells, memory B cells, plasma cells, and mast cells^58^). This overlap was significantly greater than expected by chance (1.46-fold enrichment, empirical p<0.001), supporting both the regulatory potential of these DMRs and the translational relevance of our findings. DMR-overlapping cCREs were distributed evenly across all immune cell types examined, consistent with the bulk PBMC origin of our DNA methylation data. Notably, 182 of 239 cCRE-overlapping DMRs (76%) mapped to regulatory elements shared by multiple immune cell types, including 36 (15%) present in all nine immune cell types. This indicates that gene-regulatory consequences of smoke-associated epigenetic changes may extend to various immune cell types.

To assess the functional relevance of exposure-associated DNA methylation changes, we performed gene ontology enrichment analysis of genes overlapping or within 3 kb of a DMR. Among the significantly enriched pathways (padj ≤ 0.05; Fig. 7D; Supplemental Table S8), several were relevant to the phenotypes we observed in High-WFS animals, including pathways related to oxidative stress, innate immune activation, and T cell differentiation, particularly CD8+ T cell differentiation (Fig. 7D). Inspection of individual DMR-associated genes further identified biologically relevant loci that consistently exhibited methylation differences across both sexes and/or multiple stimulation conditions (Fig. 7E). Several of these DMRs overlapped putative regulatory elements in the human genome, highlighting them as robust candidates for persistent functional epigenetic remodeling. For example, *EPHX2*, a regulator of lipid epoxide metabolism and inflammatory responses to environmental toxicants^59^, contained hypomethylated DMRs overlapping both a putative blood enhancer and an ENCODE4 cis-regulatory element in the human genome^60^. A hypermethylated intronic DMR was also found in *HIF1A*, a master regulator of hypoxic and oxidative stress responses, essential for hematopoietic stem cell maintenance^61,62^(Fig. 7E). Among DMR-associated genes involved in innate immune response, we noted *NFKBIA,* a central negative regulator of NF-κB signaling^63^, which was associated with hypermethylated DMRs in both LPS-stimulated samples of both sexes, overlapping a predicted STAT1-bound cCRE in human (Fig. 7E,F). Additional DMRs were identified at *IL12RB1*^64^, *DOCK1*^65^, and *RARRES2*^66^, genes which are implicated in inflammatory cytokine signaling, leukocyte migration, phagocytosis, and macrophage recruitment. Among DMR-containing genes implicated in CD8^+^ and T-cell differentiation, we found *TCF7,* which maintains naïve and stem-like CD8^+^ T-cell states^67^, *PRDM1,* which promotes terminal effector differentiation by antagonizing TCF7-dependent transcriptional programs^68^, *RORC*, which contributes to T-cell lineage specification and effector function^69^ (Fig. 7E).

Notably, we also observed significant enrichment of hematopoiesis-related pathways, together with DNA methylation changes in several regulators of hematopoietic stem and progenitor cell (HSPC) biology. For example, *CDKN2B*, which contributes to regulation of HSPC quiescence and self-renewal^70,71^, contained multiple hypermethylated DMRs across all three stimulation conditions in High-WFS males. These DMRs overlapped an ENCODE blood H3K27ac ChIP-seq peak and a predicted GeneHacer regulatory element interacting with the *CDKN2B* gene (Fig. 7E,F). Altered DNA methylation patterns at *CDKN2B* have been associated with environmental exposures to pesticides^72^, and genetic variation at this loci has been implicated in several human diseases^73^, highlighting it as an environmentally responsive and disease-relevant locus. We also identified DMRs associated with *FOXO3*, an essential regulator of HSC quiescence and oxidative stress resistance^74^, *RUNX1*, a master regulator of definitive hematopoiesis^75^, *BCOR*, an epigenetic regulator of HSPC fate and myeloid differentiation^76^ and *ARHGAP19*, implicated in hematopoietic progenitor proliferation and T cell biology^77^ (Fig. 7E). Collectively, these findings indicated that early-life wildfire-associated epigenetic alterations may impact pathways regulating hematopoietic development.

## DISCUSSION

Each year, millions of infants and young children are exposed to hazardous wildfire smoke during critical windows of development when environmental insults can have lasting effects on their health^3^. As wildfire smoke exposure continues to grow^4^, understanding the long-term consequences of early-life exposure and their underlying mechanisms will be essential for improving risk assessment, identifying susceptible individuals, and developing strategies to monitor, prevent, and mitigate long-term health effects. Despite a rapidly growing literature on wildfire smoke, much of our understanding of the biological mechanisms underlying smoke-related health effects remains informed by studies of tobacco smoke and urban air pollution in adults. However, wildfire smoke represents a distinct environmental exposure. In addition to its unique chemical composition, such as higher proportion of ultrafine particles and greater oxidative and inflammatory potential^78^, it also exhibits a seasonal, episodic, variable and often intense exposures lasting from days to months^79^. Likewise, findings from adults cannot be assumed to apply to pediatric exposures, when the developing body may be more susceptible and respond differently to environmental insults^80^. Together, these differences underscore the importance of studying real-world wildfire smoke exposure in pediatric models. By leveraging a cohort of adolescent rhesus macaques at ONPRC naturally exposed to wildfire smoke during infancy, we show that a single severe wildfire smoke exposure in early-life is associated with long-term attenuation of innate immune proinflammatory responses, as well as remodeling of the cytotoxic T-cell compartment. These changes were accompanied with persistent wide-spread transcriptomic and epigenetic alterations.

The persistent attenuation of LPS-induced inflammatory response we detected following early-life wildfire smoke exposure, specifically reduced IL-6 secretion in males, is consistent with previous studies in rhesus macaques experimentally exposed to ozone^16,17^ and naturally exposed to the 2008 Northern California wildfires during infancy^19^. This agreement, despite substantial differences in animal cohort and smoke exposure event highlights the robustness of this biological phenotype. By expanding our cytokine profiling relative to previous studies, we further identified reduced LTR4-induced TNF-α, GM-CSF, IL-1β, IL-10 and IFN-α responses, demonstrating that altered innate immune response extends beyond IL-6. Although males displayed stronger cytokine differences, females generally showed similar trends, suggesting that both sexes are affected, although the magnitude and nature may differ. This may reflect biological sex differences in immune regulation, including the known immunomodulatory effects of sex hormones^81^, and/or differences in developmental stage between sexes, particularly considering the earlier onset of sexual maturation in female rhesus macaques.

The attenuation of inflammatory cytokine production was further supported by transcriptomic changes in LPS-stimulated myeloid cells. Moreover, LPS-stimulated myeloid cells from High-WFS males exhibited reduced expression of *LILRA2*, an activating receptor expressed on myeloid cells^29^, and *RAP2A*, a signaling GTPase involved in LPS-induced inflammatory responses^30^. In contrast, these cells showed increased expression of *CD83* and *FSCN1*, genes associated with antigen-presenting cell maturation and acquisition of a dendritic cell-like phenotype^32^, a state specialized for myeloid cells to activate T cells and initiate adaptive immune responses. Together, these findings suggest that early-life wildfire smoke exposure does not uniformly attenuate innate immune response processes but may instead differentially affect distinct stages of myeloid activation, with evidence of attenuated inflammatory signaling alongside preserved or enhanced maturation-associated transcriptional programs. It should be noted that the transcriptional differences we detected in myeloid cells may also reflect altered gene expression within specific myeloid subsets, shifts in myeloid subclass composition, or both. Nevertheless, they are consistent with the reduced inflammatory cytokine responses observed in High-WFS animals and support persistent remodeling of innate immune function, particularly in males, following early-life wildfire smoke exposure.

In contrast to the predominantly male-derived myeloid signal, alterations within the T/NK compartment were evident in both sexes and were concentrated within cytotoxic cells. Overall, T/NK cells showed considerably more exposure-associated transcriptional changes than myeloid or B cells. Resolving this heterogeneous compartment into CD4 ^+^ and CD8^+^ subsets further amplified the transcriptional changes exclusively in CD8^+^ T cells and revealed a persistent shift from naïve toward more differentiated CD8^+^effector-memory states. Remarkably, the degree of CD8^+^ T-cell differentiation was significantly associated with the age at which animals were exposed to wildfire smoke, with earlier-life exposure associated with greater differentiation. This suggests that the timing of exposure during immune development may influence the magnitude of long-term immune remodeling. This skew appeared to reflect both a broader shift toward differentiation and expansion of a distinct CD6^+^/CD226^+^ effector cell population that was present in a subset of High-WFS animals but was nearly absent from controls. Several marker genes of this cell population, including CD6 and CD226, encode activation- or differentiation-associated T/NK receptors that have been implicated in autoimmune diseases, chronic infection, or cancer^47,50,82–86^, highlighting the potential functional importance of this phenotype. Notably, expansion and dysregulation of differentiated cytotoxic CD8^+^ T-cell populations are recurring features of chronic cigarette smoke exposure and chronic inflammatory diseases characterized by sustained immune activation^87,88^. However, unlike chronic inflammatory conditions driven by ongoing inflammation or repeated exposures, this phenotype persisted years after a single transient exposure during infancy, suggesting that early-life exposure during a critical developmental window can produce durable immune alterations. An alternative, non-mutually exclusive explanation is that early-life wildfire smoke exposure impaired host defense, leading to chronic or recurrent infections that subsequently promoted CD8^+^ T-cell differentiation. Although we cannot exclude this possibility, veterinary records from this cohort provide no evidence of chronic or recurrent infections.

Persistent DNA methylation changes are increasingly recognized as both mediators and biomarkers of the long-term biological effects of environmental exposures. These changes can reflect accelerated biological aging, alter gene regulatory programs, and may ultimately provide targets for intervention and biomarkers of disease risk. Thus, understanding how environmental exposures reshape the DNA methylome is critical for elucidating mechanisms of long-term disease susceptibility. Our DNA methylation analyses revealed a subtle genome-wide shift in DNA methylation toward an older epigenetic state, consistent with growing evidence that early-life environmental adversity and air pollution can accelerate epigenetic aging^55,89^. We also found evidence that early-life wildfire smoke exposure induces persistent epigenetic remodeling of immune regulatory networks, putative regulatory elements and immune-related transcription factor binding sites. Notably, DMR-associated genes converged on some of the same biological processes identified by our functional and transcriptomic analyses, including oxidative stress responses, innate immune signaling, CD8^+^ T-cell differentiation, and hematopoietic development. Although bulk PBMC profiling cannot identify the cellular origin of these epigenetic alterations, the enrichment of genes regulating HSPC maintenance and lineage commitment suggests that persistent remodeling of the hematopoietic compartment may also contribute to the long-term immune phenotypes observed following early-life wildfire smoke exposure.

Our study leverages a unique, naturally exposed rhesus macaque cohort to investigate the long-term consequences of a real-world wildfire smoke event, providing a highly translational model of human exposure. As with many natural exposure studies, the contributions of specific smoke constituents and co-occurring exposure-related physiological or behavioral effects (e.g., sleep disturbance^21^) could not be disentangled. Future controlled exposure studies examining smoke constituents individually will be important for identifying causal components. In addition, our sample size and cell numbers limited our ability to perform detailed analyses of less abundant immune cell subsets, particularly within the myeloid compartment. Our findings also raise the possibility that some of the observed immune alterations reflect persistent acclimation following early-life wildfire smoke exposure, although whether these changes are ultimately protective or adverse remains unknown. Lastly, while increased hospitalization during the first year following the 2020 wildfire exposure has previously been reported in the ONPRC rhesus colony^90^, whether the persistent immune alterations identified here translate into impaired host defense or other clinically relevant immune outcomes remains to be determined.

Our findings demonstrate that a single severe wildfire smoke exposure during early postnatal life is associated with persistent and extensive alterations in the immune system that remain evident into adolescence, with the greatest susceptibility occurring at the youngest ages of exposure. Our findings underscore the importance of protecting infants and young children from wildfire smoke exposure while motivating strategies to identify, monitor, and mitigate its long-term health consequences. More broadly, this study highlights early postnatal life as a critical window during which environmental exposures can durably reshape immune system development, emphasizing the need to consider childhood wildfire smoke exposures not only as transient insults, but also as potential determinants of lifelong immune health and disease susceptibility.

## METHODS

### Animal cohorts

Rhesus macaques used in this study were housed at the Oregon National Primate Research Center (ONPRC) and managed under the Institutional Animal Care and Use Committee (IACUC) approved protocol of the Oregon Health and Science University (OHSU) West Campus. All procedures in this study were approved by the OHSU Institutional Animal Care and Use Committee. Care and housing of animals was in accordance with standards established by the U.S. Federal Animal Welfare Act and the Guide for Care and Use of Laboratory Animals. During the wildfire event, all outdoor housing shelter structures were open-air without air filtration. Permanently installed sprinklers and misters provided water for vegetation and a source of cooling to minimize heat exposure.

### Exposure to the Labor Day 2020 wildfire smoke

Records of animal birth dates, as well as indoor/outdoor housing transfers dating back to September 2020, were used to retrospectively determine wildfire smoke exposure status. The early postnatal wildfire smoke-exposed cohort (High-WFS) included male and female rhesus macaques born between June 20, 2020, and September 19, 2020, that were housed outdoors September 10-19 (n = 15). The Low-WFS cohort included: (a) animals born between June 20, 2020 and September 19, 2020 that were housed in HEPA-filtered indoor facilities during the wildfire event (n = 1); (b) animals born between September 19, 2020 and February 28, 2021 with potential prenatal, but no direct postnatal, wildfire smoke exposure (n = 4); and (c) animals conceived more than one full rhesus macaque gestation period after the wildfire event (March 1, 2021-May 1, 2021), and therefore presumed to have had neither direct nor prenatal exposure to the 2020 Labor Day fires (n = 3). Animals in both exposure cohorts received comparable diets as well as veterinary care. The Oregon Department of Environmental Quality monitors a range of atmospheric pollutants across the state, including particulate matter such as PM2.5, through a network of air quality monitoring stations. The monitoring site nearest to the Oregon National Primate Research Center (ONPRC) is the Beaverton Highland Park Monitoring Station (EPA Site ID: 41-067-0111; 45.47019 N, −122.8164 W), located 7.4 km from the primate facility. Fine particulate matter concentrations were measured at this site using a Correlated Radiance Research M903 Nephelometer. For this study, the daily 24-hour average PM2.5 non-FRM/FEM mass concentration data (parameter code 88502) were obtained for the period spanning January 2019 through June 2025 to estimate ambient particulate exposure experienced by high-WFS, outdoor-housed macaques at the ONPRC. These data were retrieved from the United States Environmental Protection Agency (EPA) Air Quality System (AQS) database, which publicly provides measurements from each monitoring site for particulate matter, criteria gases, airborne toxins, meteorological conditions, and Air Quality Index (AQI) metrics.

### Blood collection and peripheral blood mononuclear cell (PBMC) isolation

High- and Low-WFS animals were approximately 3.8-4.7 years of age at the time of blood collections. All animals were considered generally healthy at time of blood collection, and no females were pregnant. Blood collections were performed opportunistically without regard to birth date or exposure status to randomize sample processing as much as possible. Rhesus macaques were sedated with 5-10mg/kg ketamine, and the venipuncture site was disinfected with alcohol or dilute chlorhexidine solution. A sterile needle with vacutainer system or collection syringe was utilized to aspirate blood. Manual pressure was applied after removal of the needle to ensure hemostasis. Monkeys are subsequently monitored until recovery from sedation. Between 8-12mL of blood was collected from each individual in Greiner Bio-One VACUETTE sodium heparin blood collection tubes (Catalog No. 456028), and blood was subsequently stored at room temperature for processing within 1 hour of collection. To extract peripheral mononuclear cells (PBMC) from whole blood, fresh blood samples were diluted 1:1 with phosphate-buffered saline (PBS) and carefully layered over a 1.5x volume Ficoll-Paque (Cytiva). Samples were centrifuged at room temperature for 45 minutes at 1700–1800 rpm without deceleration to facilitate density gradient separation. PBMCs were collected above the Ficoll layer and transferred to a new tube containing 1x PBS. Cells were then washed three times with 1x PBS. Cell counts were determined using a Countess III Automated Cell Counter (Thermo Fisher Scientific), and samples were visually inspected by microscopy to confirm cell uniformity and the absence of red blood cell contamination. PBMCs were cryopreserved in freezing medium containing 9:1 fetal bovine serum (FBS) and dimethyl sulfoxide (DMSO) and stored in a Mr. Frosty freezing container at −80 °C.

### Longitudinal weight gain analysis

Longitudinal body weight records were compiled from birth through the most recent available measurement for each animal. For most animals, weight data extended beyond the blood collection time point and continued through early 2026. All available longitudinal measurements were included to evaluate whether early postnatal wildfire smoke exposure was associated with differences in growth trajectories over time. Longitudinal body weight measurements were analyzed using linear mixed-effects models implemented in the R package nlme (v3.1). Age at each measurement was calculated as the number of days between the measurement date and date of birth. Body weight was modeled as a function of age, sex, exposure status, and all interaction terms (Weight ∼ age × sex × exposure). Animal identity was included as a random intercept to account for repeated measurements obtained from the same individual. Statistical significance of fixed effects was determined using analysis of variance of the fitted mixed-effects model. Predicted growth trajectories were plotted separately by sex and exposure group.

### *Ex vivo* immune stimulations of PBMC

From each of the 23 animals, 5-10 million frozen PBMCs were thawed in a 37°C water bath and then transferred into 10 mL of pre-warmed R10 cell culture medium (RPMI supplemented with 10% Newborn Calf Serum, L-Glutamate, Sodium Pyruvate, penicillin streptomycin and 50nM of β-Mercaptoethanol) per 1 mL of frozen cells. The cell suspension was centrifuged at 130 g for 12 minutes to gently pellet. The supernatant was aspirated, and the cell pellet was resuspended in 1 mL of R10 medium using gentle pipetting to minimize cell lysis. Cell viability and counts were assessed by mixing 10 µL of the cell suspension with 10 µL of 0.4% Trypan Blue (Thermo Fisher Scientific) and analyzing the mixture using a Countess III Automated Cell Counter (Thermo Fisher Scientific) to confirm >90% cell viability. For each sample, cells were aliquoted into three separate plate wells (1x 10^6^ - 3x 10^6^ cells/well) for immune stimulation assays. Our three stimulation conditions included: (a) incubation with media-only without any immune stimulant (i.e., unstimulated); (b) in presence of 1000 ng lipopolysaccharide (i.e., LPS); or (c) with 1000 ng phorbol 12-myristate 13-acetate (i.e., PMA), without ionomycin. As ionomycin was not included, the PMA condition is expected to induce a more moderate activation response than the commonly used PMA/ionomycin stimulation protocol, allowing us to investigate early activation signaling induced by PKC signaling. All cells were incubated at 37 °C with 5% CO_2_ overnight for 10 hours using a Tritech Digitherm Unibator. Samples were held at 4 °C at the end of the incubation period before they were centrifuged at 300 g for 10 minutes at 4 °C to pellet cells. From each well, supernatants were collected and cryopreserved at −20°C for downstream Luminex assay, while cell pellets were freshly resuspended in 1 mL of R10 medium and moved to 1.5 mL tubes for immediate transfer into single-cell RNA sequencing (scRNA-seq) assays. All surplus cells from the scRNA-seq assay were cryopreserved in freezing medium containing FBS and DMSO (9:1) and stored in liquid nitrogen until DNA methylation assay.

### Luminex quantification of immune analytes

Frozen aliquots of media supernatant from each individual immune stimulation assay (n=69) were submitted for Luminex quantification of 27 cytokines: CCL2/JE/MCP-1, CCL4/MIP-1 beta, CCL5/RANTES, CCL20/MIP-3 alpha, CD40 Ligand/TNFSF5, CXCL10/IP-10/CRG-2, CXCL11/I-TAC, CXCL13/BLC/BCA-1, GM-CSF, Granzyme B, IFN-alpha, IFN-beta, IFN-gamma, IL-1 beta/IL-1F2, IL-2, IL-4, IL-6, IL-7, IL-8/CXCL8, IL-10, IL-12 p70, IL-13, IL-17/IL-17A, IL-21, PD-L1/B7-H1, PDGF-AA, PDGF-BB, TNF-alpha, and VEGF. The Luminex assay was performed following the manufacturer’s instructions (R&D Luminex Performance NHP XL Cytokine Panel, Catalogue number: FCSTM21, R&D Systems) in the Endocrine Technologies Core (ETC) at Oregon National Primate Research Center (ONPRC). The plate was read on a Luminex-200 instrument (Luminex Corporation, Austin, TX) and analyzed with Milliplex Analyst software (MilliporeSigma, Burlington, MA) using a logistic 5P weighted curve. Sample values were adjusted for dilution as needed. Lower limit of quantification (LLOQ) was determined using the lowest curve point at least 2 SD above the blank value. Kit performance was determined using kit-included QCs run in duplicate, as well as an in-house QC of aliquoted monkey serum run in quadruplicate. To account for inter-individual differences, background-corrected cytokine responses were calculated for each analyte within each animal by subtracting unstimulated values from immune stimulated conditions (e.g., LPS - unstimulated), and these delta values were used for downstream analyses. Statistical comparisons between High- and Low-WFS animals were performed both across combined sexes and stratified by sex using Wilcoxon rank-sum tests.

### Single-cell RNA sequencing (scRNA-seq) library generation

Immediately following immune stimulation assays, PBMC samples (n=69) were assessed for cell count and viability using a Countess III Automated Cell Counter (Thermo Fisher Scientific). All 69 samples exhibited >80% cell viability and were thus used for downstream processing. scRNA-seq libraries were generated using the ScaleBio Single Cell RNA Sequencing Platform with ScalePlex sample fixation and multiplexing (ScaleBio PN 950884 and PN 1072042) following the manufacturer’s protocol. Briefly, for each condition, 2 x 10 viable cells per sample were used as input for ScalePlex fixation and sample indexing. Cells were labeled with ScalePlex oligonucleotide-conjugated antibodies according to the manufacturer’s instructions, to enable multiplexing across all 69 samples. Following labeling, the cells were washed, pooled into a single suspension, and fixed in methanol to preserve RNA integrity for downstream processing. The pooled and fixed cells were counted using a hemocytometer, and approximately 1 x 10 cells were distributed at 1 x 10 cells per well across a 96-well plate, with the remainder of the fixed cells being distributed across a ScaleBio Extended Throughput (XT) plate for the first round of combinatorial indexing. Cells were then permeabilized, and the RNA was subjected to reverse transcription using well-specific barcoded primers to capture mRNA transcripts. Cell samples were again pooled and redistributed into a 384-well plate for additional rounds of indexing via ligation, allowing for sufficient barcode complexity to uniquely label individual cells. Samples were then pooled, and cells were recounted using a hemocytometer and redistributed at approximately 1,600 cells per well into a final 96-well plate for combinatorial indexing, second-strand synthesis, and enzymatic cleanup. Sample libraries then went through tagmentation and index PCR amplification (14 cycles) to generate sequencing-ready fragments. Libraries derived from both standard and extended throughput workflows were then pooled separately, and each of the two library pools underwent two rounds of post-PCR SPRI bead cleanup (Beckman Coulter) for purification and size selection. The final, pooled scRNA library and extended throughput scRNA library were eluted, quantified using a Qubit fluorometer (Thermo Fisher Scientific), and assessed for fragment size distribution using a Bioanalyzer system. In parallel, ScalePlex sample barcode libraries were generated by PCR amplification from an aliquot of unpurified material, to enable sample demultiplexing and enrichment of hashing oligonucleotides for sequencing. A total of four pooled libraries (scRNAseq and ScalePlex libraries from both standard and extended throughput workflows) were sequenced on an Illumina NovaSeq-compatible platform using paired-end sequencing at Novogene Corporation.

### scRNA-seq data pre-processing and primary analysis

Raw BCL sequencing files were processed using the standard ScaleBio pipeline with default parameters, including default settings for ScalePlex demultiplexing, to generate gene expression count matrices. A metadata table was compiled containing animal ID, sex, wildfire smoke exposure status, immune challenge condition, and age at wildfire exposure onset. Gene count matrices and metadata were imported into Seurat v5^91^ to generate Seurat objects using default settings. Cells with ≤250 or ≥8500 detected genes were excluded, and only cells with mitochondrial transcript content <5% were retained. Cells that could not be confidently assigned to a ScalePlex well were removed from further analysis. Data were log-normalized using a scale factor of 10,000. The top 2,200 variable genes were identified and filtered to exclude genes known to produce unreliable counts or technical artifacts, including genes within the HLA class I region and gene sets previously reported to contribute to sample-to-sample batch effects, such as ribosomal genes^26^. These filtered genes were excluded only from dimensionality reduction analyses (PCA/UMAP). Data scaling and principal component analysis (PCA) were then performed. Doublets were identified using scDblFinder^92^ with an expected doublet rate of 5%, based on guidance from ScaleBio, and predicted doublets were removed. Following doublet removal, the Seurat workflow was repeated beginning from the normalization step. A small single cluster of cells with low or absent *PTPRC* expression, consistent with dead or non-immune cells, were excluded and the remaining cells were annotated into coarse immune cell types (B cells, T/NK cells, myeloid cells and unknown) using RIRA^26^. Cells receiving unknown cell classifications were excluded from downstream analyses. Cells annotated as T/NK cells were extracted, reclustered and subsequently further classified into T cell subsets (CD4^+^, CD8^+^, and gamma/delta T cells) using RIRA with default settings.

scRNA-seq-derived cell composition differences between High- and Low-WFS animals were evaluated at the individual animal level using the speckle R package with the *propeller* function^93^. Briefly, for each immune challenge condition separately (i.e., unstimulated, LPS, and PMA), proportions of major immune cell classes (B cells, myeloid cells and T/NK cells) and T cell subclasses (CD4^+^, CD8^+^ and gamma/delta T cells) identified from the scRNA-seq data were compared between wildfire smoke-exposed and unexposed animals, stratified by sex.

### Identification of differentially expressed genes (DEGs)

Differential gene expression analysis was performed using a pseudobulk strategy to account for biological replication at the animal level. Analyses were conducted separately for each major immune cell population identified by RIRA^26^, including T cells, B cells, myeloid cells, as well as CD8^+^ and CD4^+^ cells. To account for sex-specific effects observed previously and, in this study, males and females were analyzed independently. For each cell type, raw UMI counts were aggregated across all cells belonging to a given animal and stimulation condition using Seurat’s *AggregateExpression* function^91^, generating pseudobulk count matrices at the animal-by-challenge level. Samples contributing fewer than 100 cells to a given T cell, T cell subtype, B cell or 90 myeloid cells were excluded from differential gene expression analyses to ensure stable expression estimates. Differential expression testing was performed using DESeq2^94^. A combined experimental factor representing both wildfire smoke exposure status and immune stimulation condition was used in the design formula (∼ group), where group levels corresponded to exposed or unexposed animals under unstimulated, LPS or PMA conditions. This framework enabled direct contrasts between exposed and unexposed animals within each stimulation condition, as well as comparisons of stimulated versus unstimulated responses within each exposure group.

DESeq2 size-factor normalization, dispersion estimation, and Wald tests were performed using the standard workflow. Differential expression results were reported as log2 fold changes, p-values and Benjamini-Hochberg adjusted p-values. Significant differentially expressed genes identified in each comparison (padj≤ 0.05) were subsequently used for visualization and downstream pathway enrichment analyses.

### Gene signature set scoring using UCell

To compare pathway-level transcriptomic responses to LPS and PMA immune stimulation in High- and Low-WFS, we assessed inflammatory and TNF-mediated transcriptional responses using the MSigDB Hallmark Inflammatory Response and Hallmark TNFα Signaling via NF-κB gene sets^27^, each containing 200 genes. Gene symbols were matched to the rhesus macaque expression matrix by gene name, and only genes detected in the dataset were retained. To generate cell type- and stimulus-specific signatures, Hallmark gene sets were further restricted to genes significantly upregulated following LPS or PMA stimulation within each cell type (myeloid, T/NK, or B cells), based on pseudobulk differential expression analysis described above. We then used the UCell tool^28^ to calculate UCell scores for each cell using the resulting stimulus-specific gene sets. For each animal and condition, pathway activity was summarized using three complementary metrics: the fraction of cells with a non-zero UCell score, the median UCell score, and the 75th percentile (P75) UCell score across cells of the corresponding cell type. Stimulation-induced pathway responses were quantified as the change in each of these UCell score relative to unstimulated (ΔUCell = stimulated − unstimulated). Exposure effects were assessed by comparing delta values between High- and Low-WFS animals separately in males and females using two-sided Student’s t-tests.

To assess cytotoxic potential within the T/NK compartment, we calculated a cytotoxicity signature score using UCell^28^, based on canonical cytotoxic effector genes (*PRF1, GNLY, NKG7, GZMA, GZMB, GZMH, GZMK, and GZMM*). For each animal, cytotoxicity was summarized as the fraction of T/NK cells with a non-zero cytotoxicity UCell score and the median cytotoxicity UCell score across cytotoxic T/NK cells (UCell >0). These metrics were compared between exposure groups within unstimulated, LPS, and PMA conditions using the Wilcoxon rank-sum test to evaluate differences in cytotoxic potential and activation state.

### Gene ontology analysis using Gene Set Enrichment Analysis (GSEA)

Results from DEG analysis comparing High- and Low-WFS males and females from each cell type were analyzed using Gene Set Enrichment Analysis (GSEA) implemented in GSEApy (v1.1.13) in pre-ranked mode^95^. Genes were ranked from the most upregulated to the most downregulated by using the signed negative natural log-transformed p-value, incorporating both statistical significance and direction of expression change. Enrichment analysis was performed against the GO Biological Process 2025, using all genes included in the DEG analysis as the background gene set, ensuring that enrichment statistics were calculated relative to the full set of expressed genes. For each cell type, all pathways identified as significant in at least one condition (FDR q-value ≤ 0.05 and NOM ≤ p-value 0.001) were retained for downstream analysis. To identify broader functional modules, gene ontology (GO) terms were clustered based on gene overlap using pairwise Jaccard similarity calculated from pathway gene sets. A similarity network was generated using NetworkX (v3.6)^96^, where GO terms were connected if Jaccard similarity was ≥0.1, and clusters were defined as connected components within the graph. Dot plot visualizations were generated using Matplotlib (v3.10.8)^97^.

### Effector Differentiation Score (EDS) analysis and CD8^+^ Tem re-clustering

The Effector Differentiation Score (EDS) was calculated for all T/NK cells using RIRA with default parameters, as previously described^26^. Established EDS thresholds validated in both human and rhesus macaque T/NK cells were used to classify CD4+ and CD8+ T cells into naïve (T_n_; EDS ≤ 2), central memory (T_cm_; EDS 2-6), and effector memory (T_em_; EDS ≥ 6) populations. The fractions of CD8^+^ and CD4^+^ T cells classified as naïve (T_n_), central memory (Tcm), or effector memory (Tem) were compared between High- and Low-WFS animals using the Wilcoxon rank-sum test. Males and females were analyzed together, as sex-specific effects were not observed. To assess the relationship between differentiation skew and age-at-exposure, the median CD8^+^ T cell EDS was calculated for each animal. Spearman rank correlation was used to assess the correlation between median EDS of High-WFS animals and their postnatal age (in days) on Sep 9^th^ 2020. The CD8^+^ T cells classified as Tem (EDS ≥ 6) were subsetted and reanalyzed using Seurat^91^. Following normalization and dimensionality reduction, cells were clustered and visualized using UMAP. Cluster-specific marker genes were identified using Seurat’s *FindMarkers* function, and cluster identities were assigned based on the expression of these markers and their known biological functions reported in the literature.

### Genomic DNA extraction and reduced representation bisulfite sequencing (RRBS) library generation

Genomic DNA was extracted from frozen PBMCs following three immune conditions (n = 69) using the Puregene Cell DNA Extraction Kit (Qiagen Cat. 1126826), following the manufacturer’s protocol. Briefly, cryopreserved PBMC samples were thawed, transferred to 1.5 mL tubes, and centrifuged at 300 g for 10 minutes to gently pellet cells. Freezing medium was aspirated, and cell pellets were lysed using Cell Lysis Solution with thorough vortexing to ensure complete disruption. RNA was digested with RNase A, and proteins were precipitated using Protein Precipitation Solution. Samples were centrifuged at 18,000 g for 1 minute to pellet precipitated proteins, and the supernatant, containing DNA, was transferred to isopropanol for DNA precipitation. DNA was pelleted by centrifugation at 18,000 g for 2 minutes, washed twice with 70% ethanol, and subsequently air-dried. DNA pellets were then rehydrated in DNA Hydration Solution overnight at 65 °C. DNA concentration was quantified using a Qubit 1x High Sensitivity (HS) Assay (Thermo Fisher Scientific) prior to library preparation.

Reduced representation bisulfite sequencing (RRBS) libraries were prepared by digesting 100 ng of genomic DNA per with MspI to enrich for CpG-rich regions. Fragmented DNA underwent end repair, A-tailing, and ligation to methylated adapters using Ultra II DNA Library Prep modules (New England Biolabs), with methylated adapters diluted 1:10 to optimize ligation efficiency and prevent adapter dimers. Following ligation, DNA fragments were size selected to approximately 200 bp using AMPure XP magnetic beads (Beckman Coulter Cat. A63881). Adapter-ligated libraries were subject to bisulfite conversion using the EZ DNA Methylation-Gold Kit (Zymo Research Cat. D5006), which converts unmethylated cytosines to uracil while preserving methylated cytosines. Converted DNA was cleaned and desulphonated, and subsequently PCR amplified using NEBNext Q5U polymerase and NEBNext Multiplex Oligos for Illumina (New England Biolabs) to generate uniquely indexed libraries. Amplified libraries were finally purified using AMPure XP beads. The generated RRBS libraries were quantified using a Qubit HS assay (Thermo Fisher Scientific). All 69 libraries were evaluated on a Bioanalyzer system to verify fragment size distribution and absence of primer and adapter dimers prior to sequencing. Libraries were sequenced paired-end on an Illumina NovaSeq 25B lane at Novogene Corporation.

### Differential DNA methylation analysis

Sequencing quality was assessed with FastQC v0.11.9^98^, and reads were trimmed with TrimGalore v0.6.7^99^ using the -rrbs parameter. Trimmed reads were aligned to the rhesus macaque reference genome (rheMac10) with Bismark v0.24.1^100^ using default parameters. We applied the filter_non_conversion function from Bismark^100^ on the alignment bam files to remove any reads originating from inefficiently bisulfite-converted reads (based on non-CpG DNA methylation estimates) and the resulting bam files were used as input to bismark_methylation_extractor to obtain coverage files and cytosine reports for downstream use. Differential methylation analysis was conducted with DSS v2.48.0^101^ in R 4.3.3. The cytosine reports from Bismark were used as input to merge methylation data from both strands for the same CpG loci. To assess effects of smoke exposure on each sex individually, male and female samples were separated, followed by filtering for CpGs with at least 5X coverage in a majority of samples per group, where group was defined as smoke exposure_immune stimulation. This resulted in 3,758,551 CpGs for the female sample dataset, and 3,779,825 CpGs for the male sample dataset. The model was defined as ∼ group for each dataset, and the model was fit with the DMLfit.multiFactor function with smoothing set to TRUE. DML (differentially methylated loci) testing was performed with the DMLtest.multiFactor function to test the effect of smoke exposure in each stimulation (exposed vs unexposed). DMRs (differentially methylated regions) were obtained from the DML results using the callDMR function, with the following parameters: p.threshold = 0.001, minCG = 3, dis.merge = 100, minlen = 50. For each DMR, mean methylation differences between exposure groups were calculated by averaging CpG-level methylation values within each region and computing the difference between group means. DMRs were annotated to nearby and overlapping genes with ChIPseeker v1.32.0^102^ using NCBI RefSeq annotation for rheMac10.

Motif enrichment analysis was performed on distal intergenic DMRs using the findMotifsGenome.pl function from HOMER^103^ (v5.1). Gene ontology (GO) enrichment analysis was performed on genes overlapping or within 3Kb of a DMR using the EnrichR^104^ function implemented in GSEApy^95^ (v1.1.13), with the GO Biological Process 2025 gene set. The background gene set consisted of RefSeq genes annotated in the rheMac10 genome. To account for potentially opposing effects of DNA methylation changes, genes were analyzed in four independent subsets per sex: promoter hypermethylated, promoter hypomethylated, gene body hypermethylated, and gene body hypomethylated regions. Enrichment results from all subsets were subsequently combined. For pathways identified in multiple GO analyses within the same sex, the result with the lowest adjusted p-value was retained as the representative enrichment score.

To facilitate comparison with human regulatory datasets, rhesus macaque DMR coordinates were lifted over to the human hg38 genome using the UCSC LiftOver tool^105^ with default cross-species settings. A total of 1,161 of 1,435 DMRs (81%) were successfully mapped. We generated a merged reference set of candidate cis-regulatory elements (cCREs) from adult human single-cell epigenomic datasets (CATlas), including macrophages, alveolar macrophages, mast cells, naïve T cells, CD4+ T cells, CD8+ T cells, NK cells, memory B cells, and plasma cells. Lifted-over DMRs were intersected with the merged cCRE dataset using BEDTools^106^. Enrichment of DMR overlap with human immune cCREs was assessed using 1,000 random permutations in which DMR coordinates were shuffled within RRBS-covered regions of the human genome using BEDTools. An empirical p value was calculated as the proportion of permutations yielding an equal or greater number of cCRE-overlapping DMRs than observed, and fold enrichment was calculated as the observed overlap divided by the mean overlap across all permutations.

### Epigenetic clock age estimation and age-associated methylation analyses

Epigenetic age was estimated using a published blood-based rhesus macaque epigenetic clock, comprising 453 genomic regions with associated coefficients. Bismark coverage files were imported with bsseq, and coverage information was extracted for respective clock regions. For each clock region, percent methylation was computed as the ratio of methylated to total read counts across the region (perRegionTotal). Because RRBS does not guarantee coverage at every clock region in every sample, missing methylation values were imputed prior to prediction using group-specific mean imputation: a missing value was replaced by the mean methylation of that region among samples of the same exposure group, requiring at least three covered samples within the group and otherwise falling back to the overall mean across samples. Under unstimulated conditions, mean per-sample coverage was 88.3% (range 82.3-96.2%). Group-specific imputation filled the remaining gaps largely from within-group information (86.3% of missing values in the unstimulated samples), with the remainder using the population-mean fallback. Predicted epigenetic age was then obtained by multiplying each sample’s regional methylation values by the corresponding clock coefficients, summing across regions, adding the intercept, and applying the Horvath inverse age transformation (age at sexual maturity set to 5 years). Age prediction follows the clock’s published construction; the clock coefficients are applied as published. To compare epigenetic age acceleration (predicted age - chronological age) between exposure groups across all three experimental conditions, a linear mixed model was fit (predicted age ∼ chronological age + exposure status + treatment + sex + (1|animal_id)) with animal ID as a random intercept to account for repeated measures.

To further test whether smoke exposure shifts methylation toward an older epigenetic state independently of the clock, we used an external whole-blood reference dataset (184,309 loci) of known age-associated regions identified using PQLseq. Loci passing a false discovery rate threshold (q < 0.05) were retained, yielding 1,211 age-associated loci (214 gaining and 997 losing methylation with age). Methylation at these loci was extracted from the unstimulated samples (mean 91.8% coverage per sample), and missing values were imputed by group-specific mean imputation; all 1,211 loci were testable after imputation. For each locus, a linear model was fit relating methylation to chronological age, sex, and exposure status. The sign of the exposure coefficient was then compared with the sign of the reference age effect, and the proportion of loci concordant with the aging direction was tested against 0.5 with a two-sided binomial test. As a complementary, magnitude-aware check, the per-locus exposure coefficients were correlated with the reference age effects across all loci using a Spearman’s rank correlation.

### Isolation of CD34^+^ hematopoietic stem and progenitor cells (HSPC) from bone marrow

Fresh bone marrow was harvested from two femur bones per animal during necropsy and stored in cold R10 media. Samples were processed within an hour of collection, centrifuged for 10 minutes at 300rcf, followed by removal of the fat layer. Resulting pellet was resuspended in 12mL of cold PBS supplemented with 2mM EDTA and vigorously mixed for 5 minutes then spun at 830rcf for 4 minutes. The resulting pellet was resuspended in 5mL 70% isotonic percoll and underlayed beneath 7mL of 37% isotonic percoll then centrifuged at 500rcf for 20 minutes without break. Next, the buffy coat was removed, washed with HBSS, spun at 830rcf for 4 minutes, resuspended in 10mL of R10 and counted using a Horiba Petra 60C cell counter. Cells were then stained using anti-CD34 conjugated to PE (clone 561, BioLegend cat no. 343606) at 1uL per 1e7 WBC followed by labeling using Miltenyi anti-PE MACs beads (Miltenyi Biotec cat no.130-048-801) and sorting through LS columns (Miltenyi Biotec cat no. 130-042-401) using a QuadroMACS separator magnet (Miltenyi Biotec cat no.130-091-051), as per manufacturer instructions. Per sample, 1e5 cells were set aside before MACs bead labeling. After cell sorting, both CD34+ and negative fractions are spun at 300rcf for 10minutes, resuspended in R10 and counted using a Countess III cell counter. From each fraction 1e5 cells were stained, along with presorted sample, using anti-CD45 (BUV395; D058-1283; BD Biosciences; cat no. 654099) and Live/Dead Fixable NearIR (Invitrogen; cat#L34982) then incubated in the dark at 4°C for 30 minutes followed by fixation using 4% PFA. Samples were collected on a FACSymphony A5 with 10,000 events collected. Flow cell analysis was performed using FlowJo v.10.1.

## AUTHOR CONTRIBUTIONS

M. Okhovat, C. Lancioni, and L. Carbone conceived and designed the study. C. Layman, D. Morrow, K. Wheeler, P. Bergstrom performed the experimental assays. L. Carbone, C. Lancioni and S.G. Hansen provided resources and supervised staff. M. Okhovat, T. Caron, B. Davis, C. Layman, T.J. Anderson, K.N. Sterner, B. Sadoughi, N. Snyder-Mackler, G.W. McElfresh, and B.N. Bimber contributed to data analysis. K. Vigh-Conrad and M. Okhovat designed the figures. M. Okhovat wrote the manuscript, with contributions, revisions, and approval from all co-authors.

## ACKNOWLEDGEMENT

The authors thank Drs. Lisa Miller and Jonah Sacha for their insightful discussions, ideas, and guidance that helped improve this study, as well as Dr. Mary Arresto for her support with the scRNA-seq experiments. We are also grateful to Rebecca Tippner-Hedges and Dr. Shawn Chavez for generously sharing laboratory space and resources. In addition, we thank the outstanding staff at the Oregon National Primate Research Center (ONPRC), particularly Cassandra Cullins, Dr. Andrew Haertel, and Wendy Price, for their assistance. DNA methylation analyses were performed through the Knight Cardiovascular Institute (KCVI) Epigenetics Service Core at Oregon Health & Science University. The Endocrine Technologies Core (ETC) performed Luminex assays and is supported in part by NIH grant P51 OD011092 for operation of the Oregon National Primate Research Center. The research reported in this publication used computational infrastructure supported by the Office of Research Infrastructure Programs, Office of the Director, of the National Institutes of Health under Award Number S10OD034224. The content is solely the responsibility of the authors and does not necessarily represent the official views of the National Institutes of Health. This study was supported in part by pilot funding awarded to M.O. from the Friends of Doernbecher Foundation at OHSU. This work is dedicated to R and N, and to all children growing up in an era of increasing wildfire smoke exposure.

